# Novel Gurmarin-like Peptides from *Gymnema sylvestre* and their Interactions with the Sweet Taste Receptor T1R2/T1R3

**DOI:** 10.1101/2023.08.28.555239

**Authors:** Halim Maaroufi

## Abstract

*Gymnema sylvestre* (GS) is a traditional medicinal plant known for its hypoglycemic and hypolipidemic effects. Gurmarin (hereafter Gur-1) is the only known active peptide in GS. Gur-1 has a suppressive sweet taste effect in rodents but no or only a very weak effect in humans. Here, eight gurmarin-like peptides (Gur-2 to Gur-9) and their isoforms are reported in the GS transcriptome. The molecular mechanism of sweet taste suppression by Gur-1 is still largely unknown. Therefore, the complete architecture of human and mouse sweet taste receptor T1R2/T1R3 and its interaction with Gur-1 to Gur-9 were predicted by AlphaFold-Multimer (AF-M) and validated. Only Gur-1 and Gur-2 interact with the T1R2/T1R3 receptor. Indeed, Gur-1 and Gur-2 bind to the region of the cysteine-rich domain (CRD) and the transmembrane domain (TMD) of the mouse T1R2 subunit. In contrast, only Gur-2 binds to the TMD of the human T1R2 subunit. This result suggests that Gur-2 may have a suppressive sweet taste effect in humans. Furthermore, AF-M predicted that Gα-gustducin, a protein involved in sweet taste transduction, interacts with the intracellular domain of the T1R2 subunit. These results highlight an unexpected diversity of gurmarin-like peptides in GS and provide the complete predicted architecture of the human and mouse sweet taste receptor with the putative binding sites of Gur-1, Gur-2 and Gα-gustducin. Moreover, gurmarin-like peptides may serve as promising drug scaffolds for the development of antidiabetic molecules.

## Introduction

Gurmarin (in reference to “Gurmar” that is mean “sugar destroyer” in Hindi) is the only disulfide-rich peptide of 35 amino acids purified from the leaves of the Indian plant *Gymnema sylvestre* (Imoto et al., 1991; Kamei et al., 1992). Gurmarin (hereafter Gur-1) contains three disulfide bridges forming a cystine knot fold (knottin fold) that confers high resistance to proteolytic, thermal, and chemical degradation (Pallaghy et al., 1994). Gur-1 suppresses the sweet taste responses to sucrose, glucose, glycine, and saccharin in rats (Imoto et al., 1991; Yoshie et al., 1994; Miyasaka and Imoto, 1995) and mice (Ninomiya and Imoto, 1995). However, Gur-1 showed no or only a very weak suppressive sweet taste effect in humans (Imoto et al., 1991). To suppress the sweet taste response, Gur-1 is thought to interact with T1R2/T1R3 receptor in rodents (Shigemura et al., 2008; Ohkuri et al., 2009; Yasumatsu et al., 2009). The T1R2/T1R3 heterodimer, which belongs to class C G-protein-coupled receptor (GPCR), also known as seven-transmembrane (7TM) receptor, is a membrane sweet taste receptor (STR) in eukaryotes (Nelson et al., 2001). STR is expressed in taste buds. T1R2 and T1R3 monomer are also expressed in the gastrointestinal tract (Bezençon et al., 2007), pancreas (Nakagawa et al., 2009; Kyriazis et al., 2012), brain (Ren et al., 2009; Kohno et al., 2016), and testis (Mosinger et al., 2013). T1R2 and T1R3 monomer consist of an N-terminal extracellular ligand-binding (LB) domain (also called Venus flytrap module (VFTM)), a cysteine-rich domain (CRD), the seven-helical transmembrane domain (TMD), and a C-terminal intracellular domain (ICD).

A systematic review and meta-analysis have shown that *G. sylvestre* supplementation has a beneficial effect on glycemic and lipid control in patients with type-2 diabetes mellitus (Devangan et al., 2021). Recently, it has been found that Gur-1 reduces significantly sucrose intake in rodents (Rayo-Morales et al., 2023). Unfortunately, Gur-1 has no or a very weak suppressive sweet taste effect on humans. Therefore, the search for gurmarin-like peptide(s) that may be effective in humans will be helpful in the development of antidiabetic molecules. In this study, the search for the gurmarin-like peptides in the *G. sylvestre* transcriptome reveals a rich arsenal of gurmarin-like peptides. Furthermore, the predicted architecture of the human and mouse sweet taste receptor T1R2/T1R3 and their interaction with Gur-1 and Gur-2 suggest that the sweet taste suppression in rodents by Gur-1 is probably due to its interaction with the region of the CRD near the TMD of the mouse T1R2 subunit. Gurmarin-like peptides could be interesting drug scaffolds which may help to reduce the craving for sugar and thus help to regulate short-term blood glucose levels.

## Materials and methods

### Search for Gur-1-like peptides in *G. sylvestre*

TblastN algorithm (http://www.ncbi.nlm.nih.gov/blast) with Gur-1 (UniProt ID: A0A3S9JKY1 (full sequence) and P25810 (mature sequence)) as query is used to search in *de novo* transcriptome (RNA-seq) sequences of: (i) leaf, flower and developing fruits of *G. sylvestre* genotype DGS-22 deposited in NCBI Short Read Archive (SRA) (SRA accession ID: SRR5965323, SRR5965320, SRR5965321) (Kalariya et al., 2019), (ii) leaves prior to flowering of *G. sylvestre* genotype DGS-3 (SRA accession ID: SRR5965322) (Kalariya et al., 2018), and (iii) plant samples (SRA accession ID: SRR7876667) and leaf tissues (SRA accession ID: SRR3664033) of *G. sylvestre* (Ayachit et al., 2019).

### Chemical properties of gurmarin-like peptides

The chemical structures and properties of gurmarin-like peptides were investigated *in silico* using PepDraw (http://www.tulane.edu/∼biochem/WW/PepDraw/index.html) as described in Maaroufi and collaborators (Maaroufi et al., 2021).

### Phylogeny

Multiple sequences alignment are conducted by Mafft (Katoh and Toh, 2008) and phylogenetic tree is constructed by PhyML (Guindon et al., 2010).

### Residue numbering

The amino acid residues in Gur-1 to Gur-9, mouse and human T1R2 and T1R3 are numbered without the signal peptide.

### Structure prediction with AlphaFold2 and AlphaFold-Multimer

The AlphaFold2 (AF2 v2.2.2) and AlphaFold-Multimer (AF-M v2.2.2) source code was installed locally on a Linux workstation with NVIDIA Quadro RTX6000 GPU. The structure of Gur-1 has already been determined (PDB ID: 5OLL), while Gur-2 to Gur-9 peptides were predicted using AF2. While AF-M was used to predict the likely complex formation with different combinations of Gur-1 to Gur-9 peptides, T1R2 and T1R3 from mouse (UniProt ID: Q925I4 and Q925D8) and human (UniProt ID: Q8TE23 and Q7RTX0) (Jumper et al., 2021; Evans et al., 2022). AF-M is also used to study the interactions of Gur-1, Gur-2 with mouse T1R2, T1R3 and the Gα-gustducin (GNAT3) (UniProt ID: Q3V3I2).

To test the accuracy of AF-M, the brazzein peptide (UniProt ID: P56552) that has been studied experimentally for its interactions with human T1R2 and T1R3 has been used as input in AF-M with human T1R2 and T1R3.

AF2 and AF-M generate five models that are ranked based on the model confidence pLDDT score (The mean predicted Local Distance Difference Test) for AF2 and the combination of pTM+ipTM scores (pTM for monomers and ipTM for the monomers interface). Thus, pTM reports on the accuracy of prediction within each protein subunit and ipTM on the accuracy of the complex. If AF-M is unsure about an interface between the monomers, those are not docked. Each of the five predicted structure was examined. The structure figures were generated with PyMOL (http://www.pymol.org).

### Molecular dynamics simulations

In order to evaluate conformational changes of Gur-1 (PDB ID: 5OLL) and Gur-2 (3D predicted by AF2), molecular dynamics (MD) simulations were performed in GROMACS (2021.4) using the OPLS-AA/L all-atom force field (Kaminski et al., 2001) as described in (Maaroufi et al., 2021), except that Gur-1 and Gur-2 MD simulations were performed for 100 ns and in triplicate.

### Structural similarity search

DALI (http://ekhidna2.biocenter.helsinki.fi/dali/), Foldseek (https://search.foldseek.com/search) and the PDB website software (https://www.rcsb.org/search/advanced/structure) were used to search for structural similarities of Gur-1 to Gur-9, T1R2 and T1R3 in the PDB and AlphaFold databases (Van Kempen et al., 2023).

### Membrane topology prediction

Membrane T1R2 and T1R3 topology prediction was conducted by MembraneFold (https://ku.biolib.com/MembraneFold/) (Gutierrez et al., 2022).

### Detection of T1R2/T1R3 tunnels/channels

CAVER software (https://loschmidt.chemi.muni.cz/caverweb/) was used for the detection and analysis of tunnels and channels in T1R2/T1R3 structures.

### Binding affinity

The PRODIGY web server (https://wenmr.science.uu.nl/prodigy/) was used to calculate the binding free energies (ΔG) of interacting protein-protein and peptide-protein complexes (Xue et al., 2016).

### Amino acids interactions in protein-protein complex

PyMOL and ContPro (http://procarb.org/contpro/index.html are used to determine the bonds between protein-protein and peptide-protein complexes.

## Results and discussion

### *G. sylvestre* transcriptome is rich in gurmarin-like peptides

Gurmarin (hereafter Gur-1) is the only active peptide extracted from the leaves of *Gymnema sylvestre* (Imoto et al., 1991; Kamei et al., 1992). In the aim to find homologs of Gur-1 (UniProt ID: P25810), a TblastN was performed using the transcriptome (RNA-seq) sequences of leaves, leaves prior to flowering, flowers and developing fruits of *G. sylvestre* deposited in the NCBI Short Read Archive (SRA) (Kalariya et al., 2018, 2019; Ayachit et al., 2019). From this search, 57 gurmarin-like peptides were found. Multiple sequence alignment (MSA) of these peptides revealed nine groups, named Gur-I to Gur-IX (Fig. S1). Gur-I group is rich in isoforms of Gur-1. Interestingly, Trp29 whose mutation to alanine completely abolishes the sweet-suppressing properties of Gur-1, is conserved in isoforms of Gur-I and Gur-II groups (Sigoillot et al., 2018). It has been found that Gur-1 has no or only a very weak suppressive sweet taste effect in humans (Imoto et al., 1991; Miyasaka and Imoto, 1995). Therefore, this arsenal of isoforms might have a more potent sweet taste inhibitory effect in humans. It is noteworthy that some of these isoforms have mutations in Cys10 or Cys17 and Cys10/Cys17. Gur-1 has three disulfide bridges (Cys3-Cys18, Cys10-Cys17 and Cys23-Cys33). It has been shown that mutagenesis of these six cysteines alters the function of Gur-1. Indeed, mutant Gur-1 analogs with only two native disulfide bonds suppressed the response to sucrose, but not those to glucose, fructose, saccharin, or glycine in rats (Ota et al., 1998b).

For the continuation of this work, one representative sequence from each group of gurmarin-like peptides was selected and named Gur-1 (from group I) to Gur-9 (from group IX) (Fig. 1). The analysis of Gur-1 to Gur-9 showed that they have different physicochemical properties (Table S1). Gur-2 is more similar to Gur-1 (48.57% of identity and 65.71% of similarity in amino acid residues). Like Gur-1, Gur-2 has a hydrophobic and hydrophilic cluster on the surface of the molecule (Ota et al., 1998a). Gur-2 is more hydrophobic than Gur-1 (Table S1). For Gur-3 to Gur-9, BlastP (expect threshold of 0.05) against the NCBI database showed no matches, indicating that they are novel knottin (cystine knot fold) peptides.

**Fig. 1.**
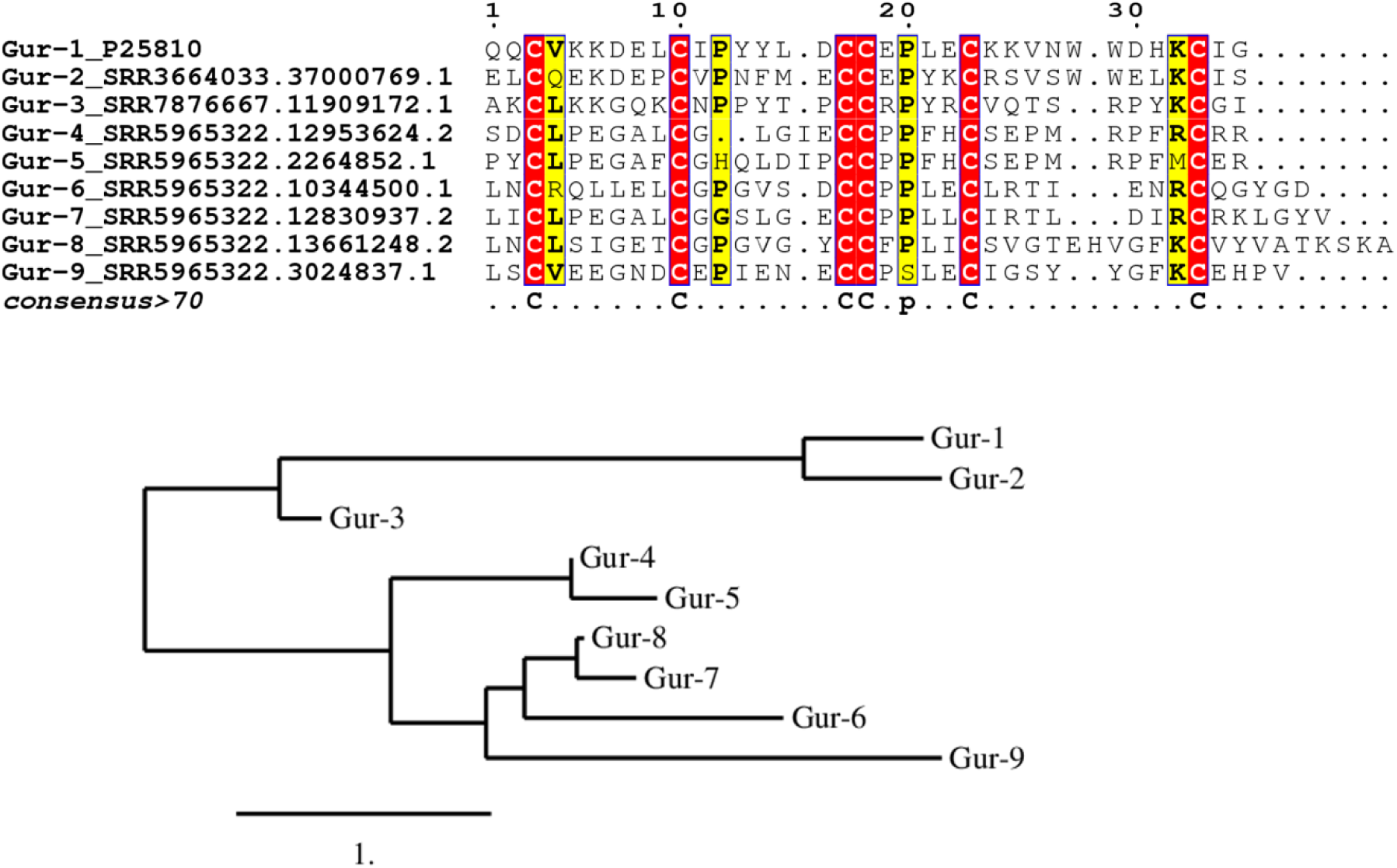
Multiple sequence alignment and clustering of Gur-1 (UniProt ID: P25810) to Gur-9 peptides. Each peptide has six conserved cysteines that are predicted to form a knottin three disulfide bonds. SRA accession ID is indicated with each sequence.

### Prediction of the structure of gurmarin-like peptides

The structure of Gur-1 is known (PDB ID: 5OLL). AlphaFold 2 (AF2) is used to predict the structures of Gur-2 to Gur-9. AF2 produces five ranked models. Each of the five predicted structures was examined. AF2 predicted structures of all gurmarin-like peptides in the except of Gur-6. Of note, in the predicted structure of Gur-5 (pLDDT score of 56.77 is bad) the three disulfide bridges are different from those in Gur-1. Therefore, ESMFold software (based on large language models) that uses only the amino acid sequence to predict the structure is used for Gur-5 and Gur-6 (Lin et al., 2023). The results of the structure alignments of Gur-1 to Gur-9 using DALI (Holm, 2020) are shown in Table S2. The structural similarity dendrogram of Gur-1 to Gur-9 shows that the structures of Gur-2 to Gur-5 are close to Gur-1, whereas Gur-6/Gur-8 and Gur-7/Gur-9 are distant from Gur-1 (Fig. S2).

### Prediction of the structure of the T1R2/T1R3 heterodimer

Subunits T1R2 and T1R3 of sweet taste receptor consist of a signal peptide, an N-terminal extracellular domain, the seven-helical transmembrane domain (TMD), and a C-terminal intracellular domain (ICD). There is no experimental structure of the monomers or the complete sweet taste receptor T1R2/T1R3. Only the extracellular ligand-binding domains (LBDs, also named Venus flytrap module (VFTM)) of the medaka fish T1R2/T1R3 heterodimer have been determined (PDB ID: 5X2M). Therefore, AF2 was used to predict the structure of the human T1R2 and T1R3 (hT1R2 and hT1R3) and the mouse T1R2 and T1R3 (mT1R2 and mT1R3) monomers (Fig. 2), and AlphaFold-Multimer (AF-M) to predict the structure of the hT1R2/T1R3 and mT1R2/T1R3 heterodimer (Fig. 4). AF-M generates five ranked models. Each of the five predicted structures was examined. The pLDDT (the mean predicted Local Distance Difference Test) score of mT1R2, mT1R3, hT1R2 and hT1R3 monomers are 84.41, 86.97, 85.81 and 87.1, respectively, indicating a high 3D models quality. While the best 3D model of mT1R2/T1R3 and hT1R2/T1R3 has a pTM+ipTM (overall predicted TM (pTM) + predicted interface (ipTM)) score of 0.68. When the ipTM score is greater than 0.6, it indicates that the protein interface is well predicted (Evans et al., 2022). The twilight zone for a good confidence of the predicted interface is found between 0.5 and 0.6. Recently, at CASP15, the accuracy of AF-M has been demonstrated in modeling the quaternary structure of proteins in comparison to bioinformatics methods used prior to AF software (Ozden et al., 2023).

**Fig. 2.**
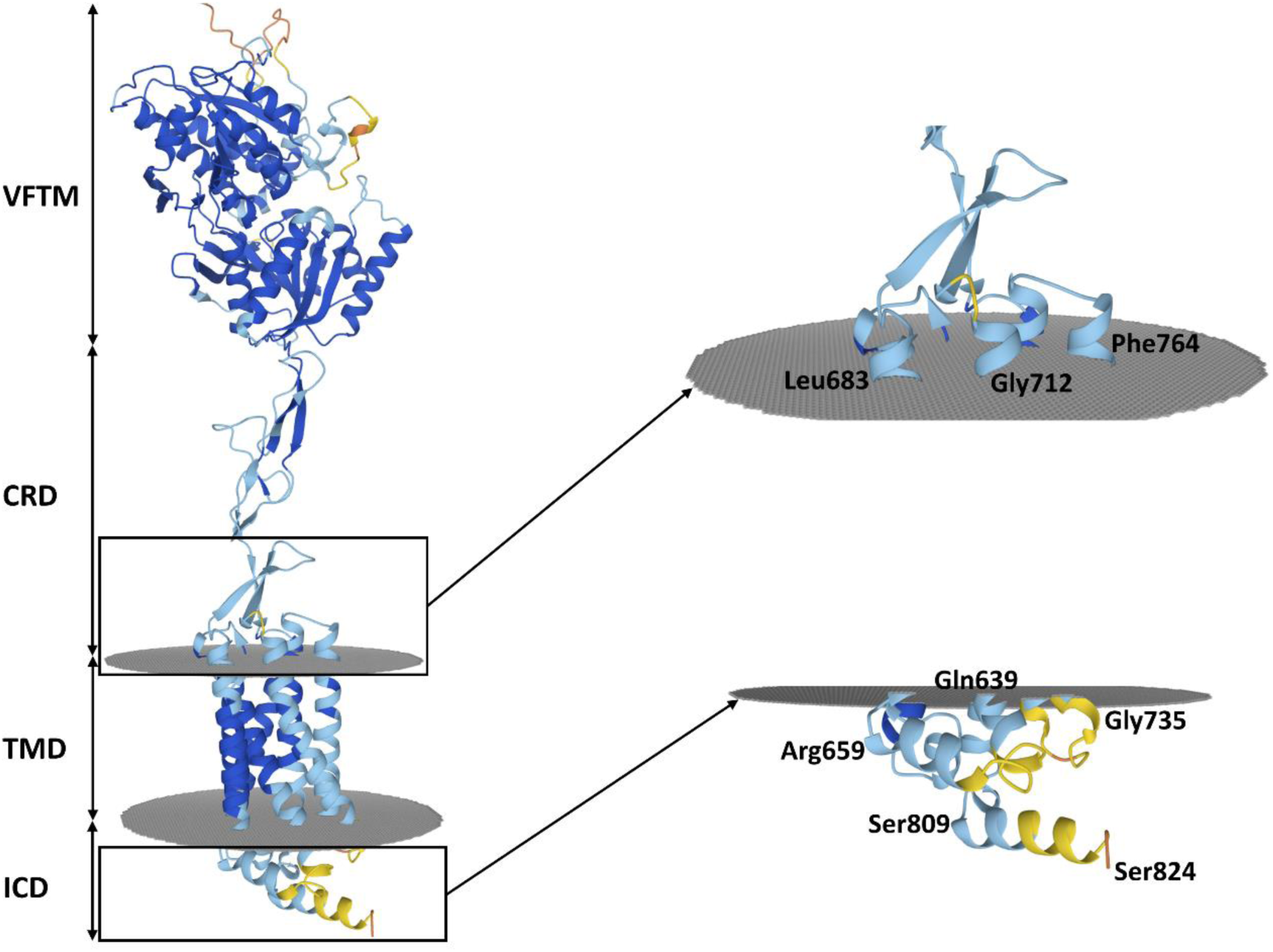
The membrane topology of the predicted structure of the mouse T1R2 (mT1R2) by AF2. mT1R2 is formed with an N-terminal Venus flytrap module (VFTM), a cysteine-rich domain (CRD), a seven-helical transmembrane domain (TMD) and an intracellular domain (ICD). The accuracy of the predicted model is visualized by different colors (pLDDT ≥ 90, 90>pLDDT≥70, 70>pLDDT≥50, 50>pLDDT). Amino acid residues of mT1R2 are numbered without the signal peptide.

Analysis of the hT1R2/T1R3 complex revealed that amino acid residues of hT1R2 involved in interactions with hT1R3 according to Park et collaborators (Park et al., 2019) are effectively located at the interface between hT1R2 and hT1R3 in the model predicted by AF-M (Fig. 3). In addition, an ionic bond between Arg217 of hT1R2 and Glu217 of hT1R3 in the predicted model is consistent with the study of Cui and coworkers who, by mutating hT1R2 Arg217 to alanine suggested that Arg217 is involved in the hT1R2/T1R3 interaction (Cui et al., 2006). The predicted binding affinity of mT1R2 and mT1R3 in the heterodimer (ΔG ⁓ –23.1 kcal/mol) is similar to that of hT1R2 and hT1R3 (ΔG ⁓ –21.0 kcal/mol).

**Fig. 3.**
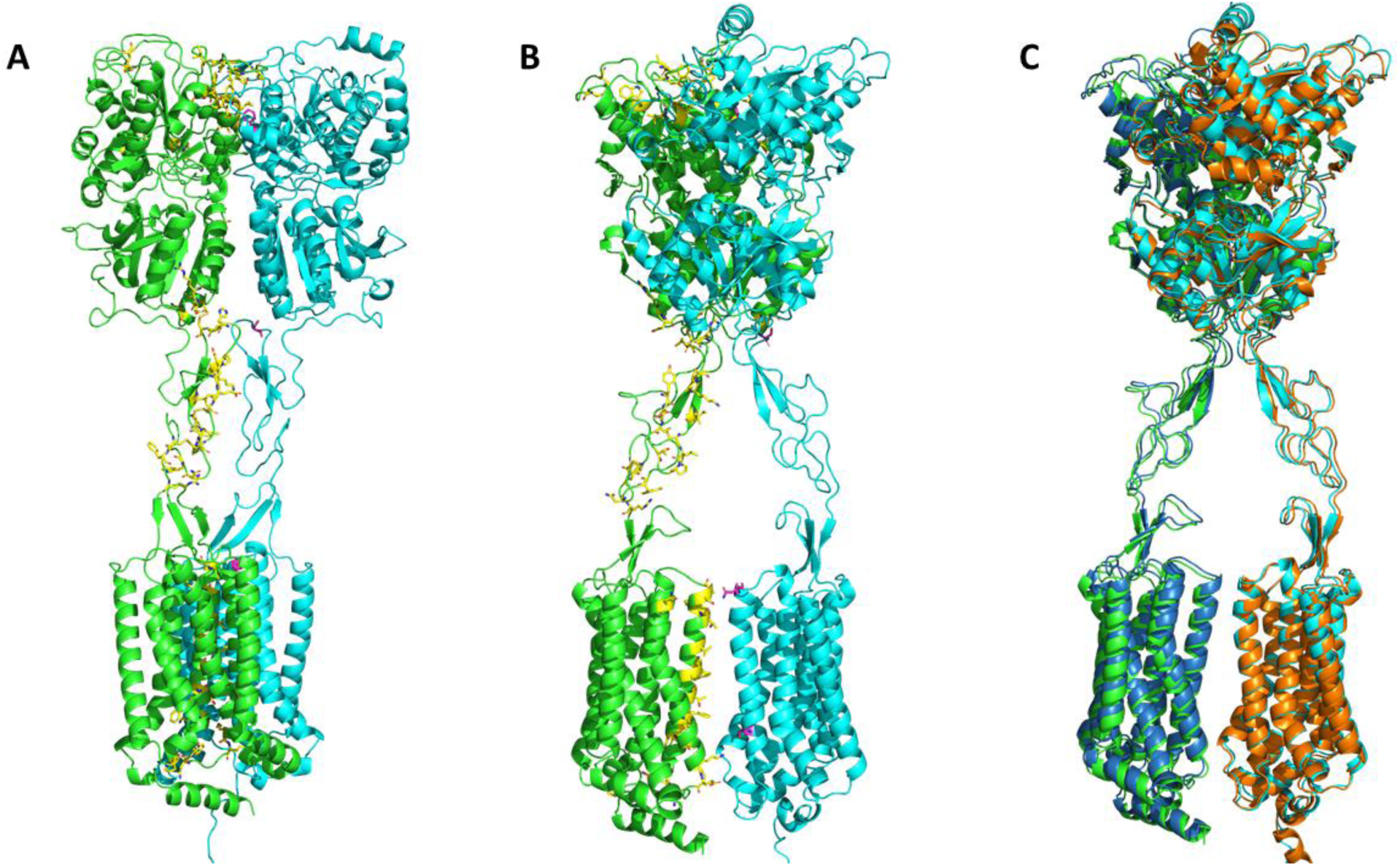
Predicted structures of human T1R2/T1R3 (hT1R2/T1R3) and mouse T1R2/T1R3 (mT1R2/T1R3) heterodimer. **A** and **B**, hT1R2/T1R3 (green/cyan) with amino acids (yellow sticks) of hT1R2 implicated in the interactions with hT1R3 according to Park et coworkers (Park et al., 2019). **C**, superposition of hT1R2/T1R3 (green/cyan) and mT1R2/T1R3 (blue/orange). The structure of mT1R2/T1R3 aligns on hT1R2/T1R3 with an RMSD of 1.84 Å.

The structure similarity search (https://www.rcsb.org/search/advanced/structure) in PDB database revealed that the predicted structure of the hT1R2 and mT1R3 monomers are structurally similar to the human calcium-sensing receptor (CaSR) (PDB ID: 7M3G_A) and human metabotropic glutamate receptor 2 (mGlu2) (PDB ID: 8JD3_A). Like T1R2 and T1R3, CaSR and mGlu2 are a class C G protein-coupled receptors (GPCRs).

Pairwise structure alignment (https://www.rcsb.org/alignment) using the jFATCAT (flexible) option, aligned T1R2 and T1R3 monomers to CaSR and mGlu2 with an RMSD of 3.1-3.21 Å at 723-765 amino acid residues, and 2.71-3.76 Å at 700-740 amino acid residues, respectively. Whereas T1R2 and T1R3 are aligned to the LBDs of the medaka fish T1R2/T1R3 heterodimer (PDB ID: 5X2M) with an RMSD of 1.89-2.18 Å on 427-440 amino acid residues.

### Prediction of interactions between gurmarin-like peptides and T1R2 and T1R3

#### Brazzein and hT1R2/T1R3 interactions as control of AF-M accuracy

It has been demonstrated that AF-M is capable of predicting peptide-protein complex with high accuracy (Chang and Perez, 2023; Teufel et al., 2023). This accuracy sometimes depends on type of proteins. Therefore, the published experimental results of the binding of Brazzein (Brz), a sweet-tasting peptide of 54 amino acids stabilized by four disulfide bonds, with hT1R2 and hT1R3 were used to test the accuracy of the AF-M software for predicting the peptide-protein interactions.

AF-M showed that Brz interacts with the region Ile516-Glu525 in the CRD of hT1R3 subunit of T1R2/T1R3 heterodimer (iptm+ptm = 0.6168) (Fig. S3). This region is involved in the responsiveness to Brz. Indeed, the functional mutagenesis analysis revealed that Ala517 and Phe520, located in the region Ile516-Glu525, jointly determine the different sweet taste sensitivity between human (taster) and mouse (non-taster) toward Brz (Jiang et al., 2004a). In the predicted hT1R2/T1R3 structure, Ala517 and Phe520 interact with two adjacent aromatic residues Tyr39 and Phe38 of Brz, respectively. Interestingly, also two adjacent aromatic residues Trp28 and Trp29 in Gur-1 are important for the sweet-suppressing properties of Gur-1 (Sigoillot et al., 2018). In addition, as for Trp29 in Gur-1 (its mutation to alanine abolishes the suppressive effect of Gur-1), the second aromatic residue Tyr39 is more critical for the Brz activity (Jiang et al., 2004; Sigoillot et al., 2018). Moreover, Ala19 of Brz binds Arg490 of hT1R3. The site-directed mutagenesis of Ala19 has demonstrated its importance for the sweetness effect of Brz (Ghanavatian et al., 2016). Therefore, these experimental results support the accuracy of AF-M for predicting peptide-protein interactions of type peptide-sweet taste receptor.

#### Interactions of gurmarin-like peptides and T1R2 and T1R3

*In situ* studies suggest that Gur-1 sensitivity is related to the coupling of T1R2/T1R3 to Gα-gustducin (Shigemura et al., 2008; Ohkuri et al., 2009; Yasumatsu et al., 2009). In addition, the recombinant Gur-1 has been shown to be responsible for the complete blockade of saccharin responses in rat T1R2/T1R3-expressing cells (Sigoillot et al., 2012a). Therefore, in the aim to understanding the molecular mechanism of sweet taste suppression by Gur-1 and gurmarin-like peptides, AF-M is used to predict the interactions of Gur-1 to Gur-9 with hT1R2/T1R3 and mT1R2/T1R3. AF-M is used with these combinations of human and mouse amino acid sequences (Gur, mT1R2), (Gur, mT1R3), (Gur, mT1R2, mT1R3), (Gur, hT1R2), (Gur, hT1R3), and (Gur, hT1R2, hT1R3) (Table 2). The interactions of Gur-1 and gurmarin-like peptides with T1R2 and T1R3 monomers are studied in the case where, in the taste cell, Gur-1 and Gur-2 bind T1R2 and/or T1R3 before the dimerization of the sweet taste receptor.

The results showed that only Gur-1 and Gur-2 interact with the T1R2/T1R3 receptor. No interactions of Gur-3 to Gur-9 are obtained with monomers nor T1R2/T1R3 heterodimer. This suggests that Gur-3 to Gur-9 may have function other than sweet taste suppression. AF-M produces five ranked models. In the case of T1R2/T1R3-Gur-1 or T1R2/T1R3-Gur-2 complexes, two models with different binding sites for Gur-1 or Gur-2 were obtained with a high pTM+ipTM score. Indeed, AF-M predicted that Gur-1 and Gur-2 interact with the extracellular and intracellular (less probable because Gur-1 and Gur-2 must cross the taste cell membrane) domains of T1R2/T1R3. Previous studies have hypothesized that Gur-1 may interact with the extracellular region (VFTM and/or CRD) of mT1R2/T1R3 but not with the intracellular region (Sigoillot et al., 2012b, 2018).

In the extracellular region, Gur-1 and Gur-2 bind to the CRD near the TMD of mT1R2 subunit of mT1R2/T1R3 with binding affinity of –7.1 and –7.7 Kcal/mol, respectively (Fig. 4 and Fig. S5). Whereas in hT1R2/T1R3 only Trp29 of Gur-2 forms a cation-pi bond with Arg706 of TMD of hT1R2 subunit with binding affinity of –5.1 Kcal/mol (Table 1, Tables S3 to S6). Interestingly, Gur-1 do not interact with hT1R2/T1R3. This is consistent with works that showed that Gur-1 has no or only weak suppressive effect in human (Imoto et al., 1991; Ninomiya and Imoto, 1995).

**Fig. 4.**
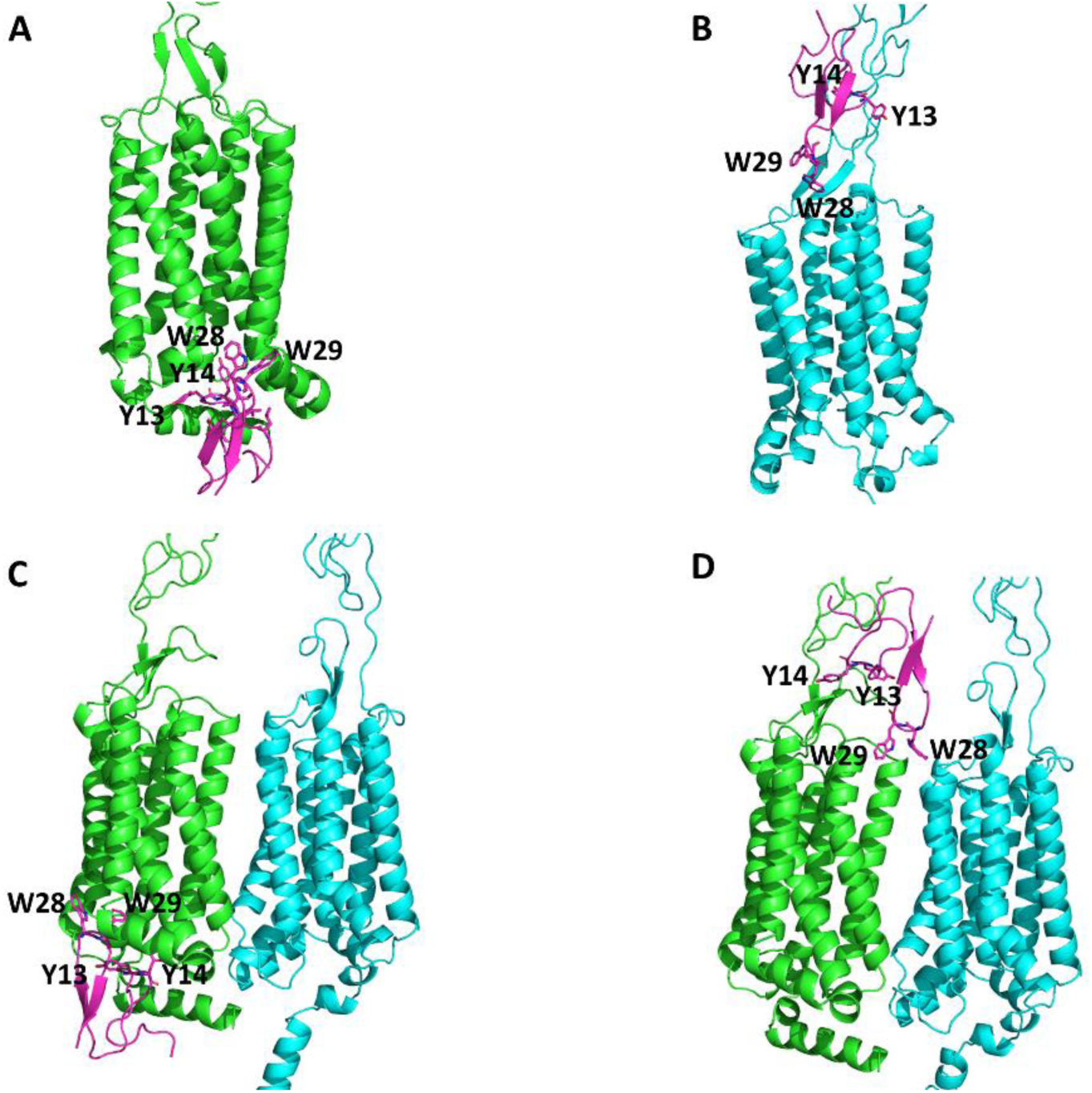
Interactions of Gur-1 with mouse T1R2, T1R3 monomers and T1R2/T1R3 heterodimer. (**A**) Gur-1 (pink) binds in the intracellular domain (ICD) of the T1R2 monomer (green) with Y13, Y14, W28 and W29 amino acid residues. (**B**) Gur-1 binds in the cysteine-rich domain (CRD) near the transmembrane domain (TMD) of the T1R3 monomer (cyan). (**C**) and (**D**) Gur-1 binds in the ICD and the CRD near TMD of the T1R2 subunit of T1R2/T1R3 heterodimer (green/cyan), respectively. For details of bonds between amino acid residues, see Table S3.

**Table 1.**
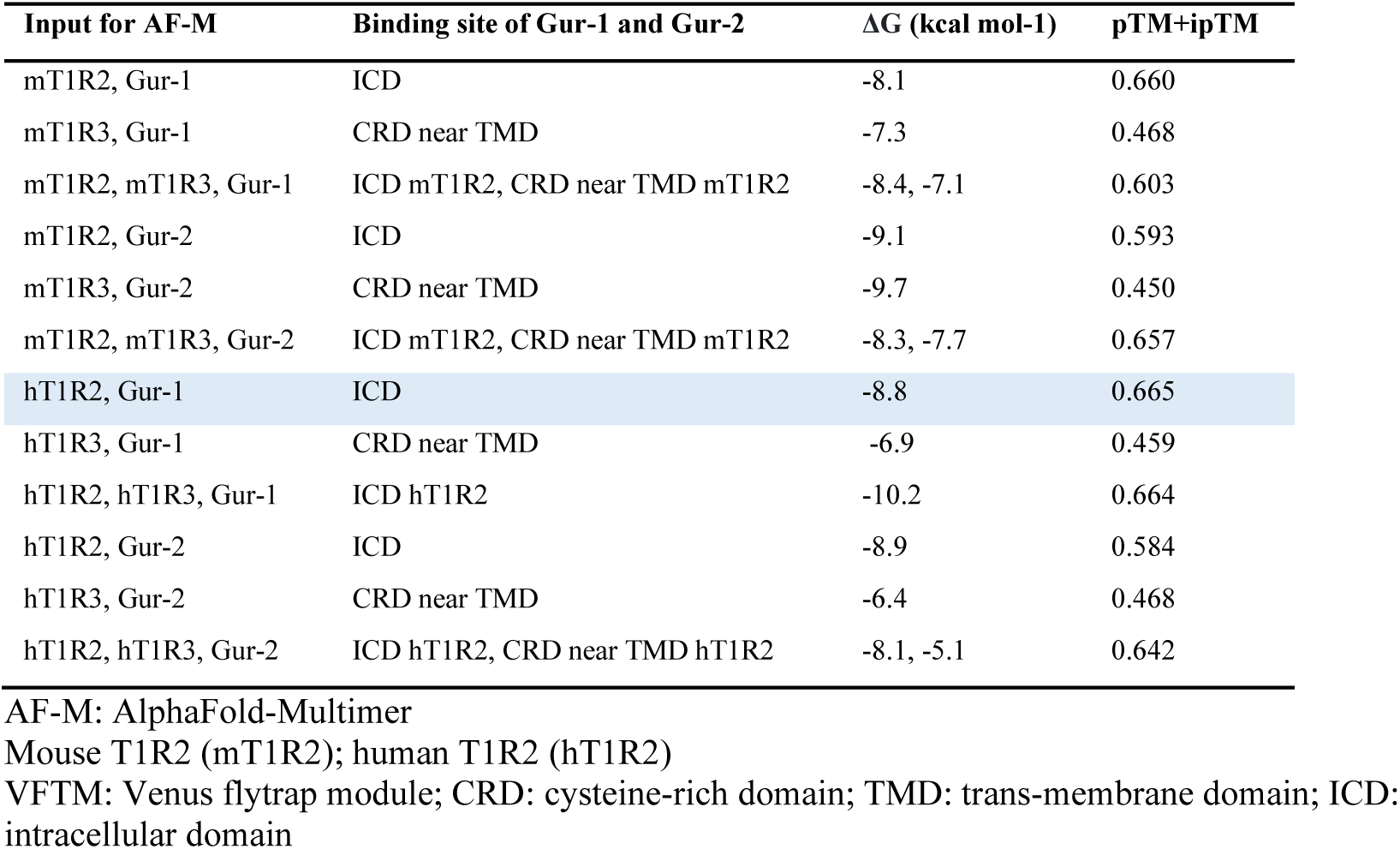
Predicted interactions by AF-M of Gur-1, Gur-2, T1R2 and T1R3.

| Input for AF-M | Binding site of Gur-1 and Gur-2 | $\Delta G$ (kcal mol<sup>-1</sup>) | pTM+ipTM |
| --- | --- | --- | --- |
| mT1R2, Gur-1 | ICD | -8.1 | 0.660 |
| mT1R3, Gur-1 | CRD near TMD | -7.3 | 0.468 |
| mT1R2, mT1R3, Gur-1 | ICD mT1R2, CRD near TMD mT1R2 | -8.4, -7.1 | 0.603 |
| mT1R2, Gur-2 | ICD | -9.1 | 0.593 |
| mT1R3, Gur-2 | CRD near TMD | -9.7 | 0.450 |
| mT1R2, mT1R3, Gur-2 | ICD mT1R2, CRD near TMD mT1R2 | -8.3, -7.7 | 0.657 |
| hT1R2, Gur-1 | ICD | -8.8 | 0.665 |
| hT1R3, Gur-1 | CRD near TMD | -6.9 | 0.459 |
| hT1R2, hT1R3, Gur-1 | ICD hT1R2 | -10.2 | 0.664 |
| hT1R2, Gur-2 | ICD | -8.9 | 0.584 |
| hT1R3, Gur-2 | CRD near TMD | -6.4 | 0.468 |
| hT1R2, hT1R3, Gur-2 | ICD hT1R2, CRD near TMD hT1R2 | -8.1, -5.1 | 0.642 |

AF-M: AlphaFold-Multimer
Mouse T1R2 (mT1R2); human T1R2 (hT1R2)
VFTM: Venus flytrap module; CRD: cysteine-rich domain; TMD: trans-membrane domain; ICD: intracellular domain

Gur-1 binds through its hydrophobic cluster (Tyr13-Tyr14 and Trp28-Trp29) to the region of CRD near TMD (Fig. 4D). It has been reported that tryptophan residues and their position in peptides play an important role in the interaction with lipid membranes (Rydberg et al., 2014). In addition, Obi and Natesan revealed the role of membrane lipids in the binding affinity, the stability of ligands at the binding site in the G-protein-coupled receptor (Obi and Natesan, 2022). They suggest considering the ligand-lipids interactions for calculating the binding affinity. Considering lipids, Gur-1 may have more affinity for mT1R2/T1R3 than its predicted binding affinity (Table 1). Therefore, the probable interaction of Gur-1 with lipids may explain why the effect of Gur-1 is not immediate, requiring 5 min to achieve maximal suppression of sweet responses (Ninomiya et al., 1999). It may also explain the long-lasting suppressive effect of Gur-1 (>2-3 h) (Imoto et al., 1991) and the ineffectiveness of the rinsing procedures (Sigoillot et al., 2018), in addition to the fact that the suppressed sucrose response after Gur-1 recovered to some extent after rinsing the tongue with 15 mM β-CD (β-cyclodextrin) for 10 min (Ninomiya et al., 1998). It has been suggested that β-CD interacts with tyrosine and tryptophan residues of Gur-1 hydrophobic cluster that facilitates the dissociation of the mT1R2/T1R3-Gur-1 complex (Imoto et al., 2001). Indeed, in this work Gur-1 and Gur-2 bind to T1R2/T1R3 through their hydrophobic cluster Tyr13-Tyr14/Trp28-Trp29 and Phe14/Trp28-Trp29, respectively.

In the cytoplasmic region (less probable because Gur-1 must cross the taste cell membrane), Gur-1 and Gur-2 bind to the intracellular domain (ICD) of hT1R2 and mT1R2 monomers as well as the T1R2 subunit of the T1R2/T1R3 heterodimer (Fig. 4A and C) with binding affinities between –8.1 and –10.2 kcal/mol (Table 1). The electrostatic surface potential showed that the ICD forms a crevice that accommodates Gur-1 and Gur-2 (Fig. S4). The interaction with the ICD of T1R2 requires that Gur-1 and Gur-2 cross the taste cell membrane. Otherwise, the results showing that Gur-1 and Gur-2 interact with the ICD have no biological value. As hypotheses, Gur-1 and Gur-2 could have a propriety (cell-penetrating peptides) to cross the membrane (Gori et al., 2023) and/or Gur-1 can enter the cell via the predicted channel between T1R2 and T1R3 subunits (Fig. S6).

Finally, Gur-1 and Gur-2 bind to the CRD near TMD in the human and mouse T1R3 monomers but not in T1R2/T1R3 heterodimer (Table 1 and Fig. 4B).

The amino acids of Gur-1 and Gur-2 involved in interactions with T1R2 and T1R3 are mainly Tyr13-Tyr14 and Trp28-Trp29 for Gur-1 and Phe14 and Trp28-Trp29 for Gur-2 (Tables S3-S6). Interestingly, hydrophobic amino acids Tyr14, Trp28 and Trp29 are showed involved in the sweet-suppressing properties of Gur-1 (Ota et al., 1998a; Sigoillot et al., 2018).

#### Interaction of Gur-2 with chimeric T1R2/T1R3 heterodimer

Gur-2 binds to the CRD-TMD region of mT1R2 subunit of mT1R2/hT1R3 with an affinity of –7.7 Kcal/mol (Fig. 5A). Whereas only Trp29 of Gur-2 interacts with Arg706 in the TMD of hT1R2 subunit with an affinity of –5.1 Kcal/mol (Fig. 5B). In the aim to empower the binding of Gur-2 to the hT1R2 subunit the region between amino acids Thr686 and Gln702 of hT1R2 is replaced by the corresponding region of mT1R2 (Fig. 5E). This replacement improves the binding affinity (–6.8 Kcal/mol) of Gur-2 to chimeric mouse hT1R2/hT1R3 comparatively to hT1R2/hT1R3 (–5.1 Kcal/mol) (Fig. 5C).

**Fig. 5.**
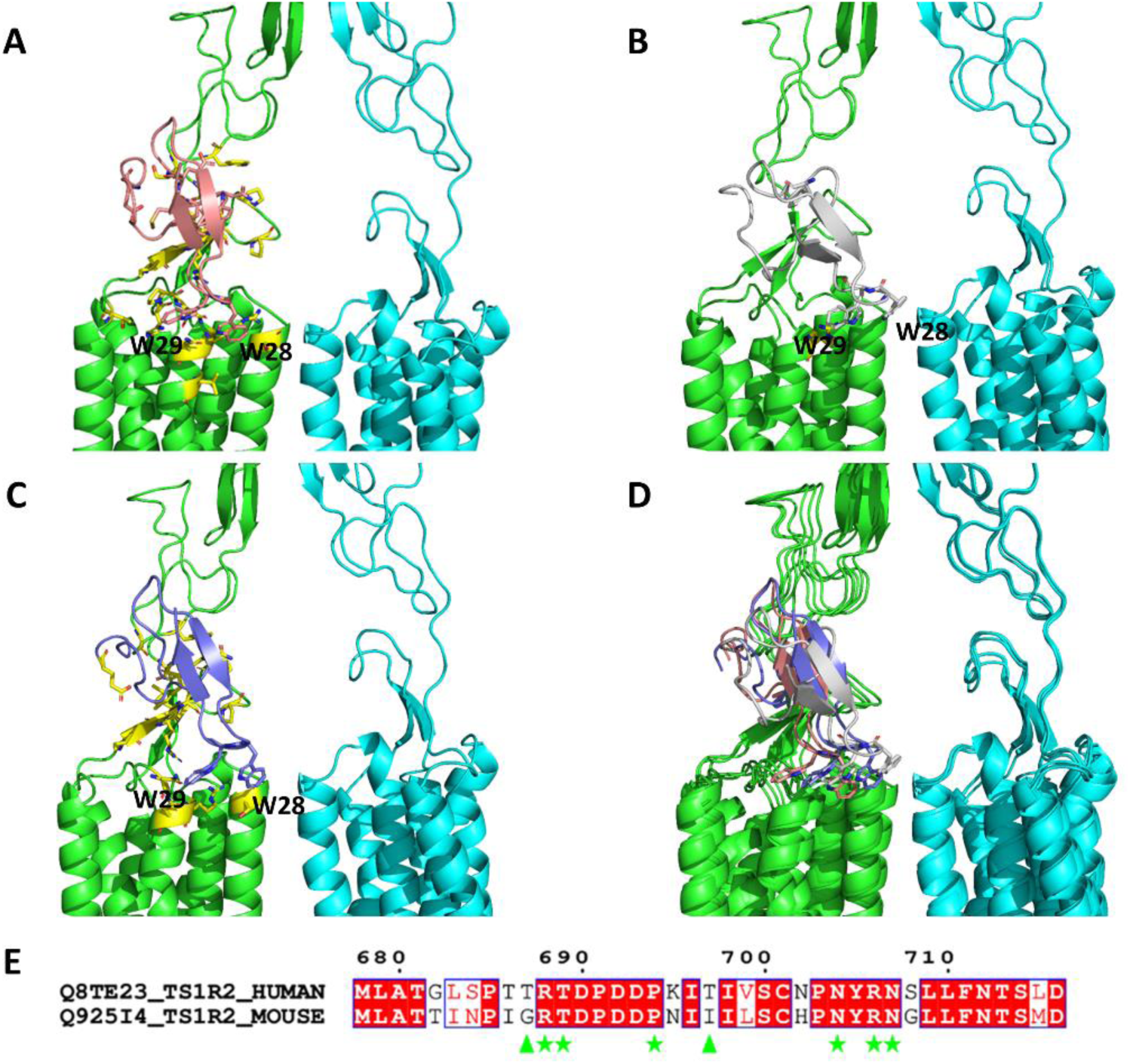
Interactions of Gur-2 with the mouse, human and chimeric T1R2/T1R3 heterodimer. (**A**) Large interaction interface (yellow) of Gur-2 (salmon) and the mouse T1R2 subunit (mT1R2). (**B**) Only the residue W29 of Gur-2 (grey) interacts with R706 of the human T1R2 subunit (hT1R2). (**C**) Large interaction interface (yellow) of Gur-2 (blue) and the chimeric T1R2/hT1R3 (The region between Thr686 and Gln702 of hT1R2 is replaced by the corresponding region of mT1R2 (**E**)). (**D**) superposition of mT1R2/T1R3-Gur-2 (Gur-2 in salmon), hT1R2/T1R3-Gur-2 (grey) and chimeric T1R2/T1R3-Gur-2 (blue). T1R2 subunit (green) and T1R3 subunit (cyan). (**E**) Pairwise alignment of the region where Gur-2 interacts with mT1R2. Amino acids in mT1R2 that interact with Gur-2 are underlined with green stars and head arrows. The latter represents the two amino acids different in mT1R2 and hT1R2.

#### Interactions of Gur-1 and Gur-2 with homodimers of T1R2 and T1R3

The use of AF-M with two sequences of mT1R2 and one sequence of Gur-1 or Gur-2 showed only the formation of mT1R2/T1R2 homodimers, but not mT1R2-Gur-1 or mT1R2-Gur-2 complexes. The same result was obtained with two sequences of mT1R3 and one sequence of Gur-1 or Gur-2. It has been suggested that the putative homodimerization of T1R3 is not functional (Shimizu et al., 2014). Recently, Belloir and coworkers found that the overexpressed hT1R2 is predominantly a dimer (Belloir et al., 2021). AF-M results indicate that Gur-1 and Gur-2 only interact with the T1R2/T1R3 heterodimer. This is consistent with the requirement of heterodimerization of T1R2 and T1R3 for sweet taste perception.

#### *In silico* competition assays between Gur-1 and Gur-2

To compare the binding affinity of Gur-1 and Gur-2 to T1R2, T1R3 and T1R2/T1R3, *in silico* competition assays were conducted (Chang and Perez, 2023; Banerjee et al., 2023). To remember that interactions of Gur-1 and Gur-2 with the ICD are considered less probable because Gur-1 and Gur-2 must cross the taste cell membrane.

For T1R2 as monomer or in T1R2/T1R3 heterodimer, Gur-1 binds or clashes with the ICD while Gur-2 binds to the CRD near TMD region (Table S7). Whereas for T1R3 monomer, Gur-1 binds or clashes with the TMD near ICD while Gur-2 binds to the CRD near TMD region. Therefore, comparatively to the results obtained with single gurmarin peptide (Table 1), Gur-2 has more affinity than Gur-1 for the CRD near TMD region of T1R2 or T1R3. That would be interesting to use Gur-1 and Gur-2 (and their isoforms) together in the experiments to know if they may have a synergistic effect on the suppression of sweet taste in humans.

#### Effect of Gur-1 or Gur-2 binding on the conformational changes

The pairwise structural alignment was performed to know whether the binding of Gur-1 or Gur-2 to the T1R2/T1R3 heterodimer changes the conformation of the complex. For mice, structural alignment of the mT1R2/T1R3 with the mT1R2/T1R3-Gur-1 or mT1R2/T1R3-Gur-2 complex showed an RMSD of 3.02 Å and 3.35 Å over 1624 amino acid residues, respectively. Whereas hT1R2/T1R3 with hT1R2/T1R3-Gur-2 complex is 3.10 Å over 1600 residues. These results show that the binding of Gur-1 or Gur-2 to the T1R2/T1R3 induce an important conformational change that may destabilize the active conformation of T1R2/T1R3 and thus prevent the transmission of the sweet taste signal from taste cell surface to intracellular signaling pathways (Perez-Aguilar et al., 2019).

### Molecular dynamics simulation

Only Gur-1 and Gur-2 among the gurmarin-like peptides interact with the sweet taste receptor. They share 51% of identity in amino acids and are structurally aligned with an RMSD of 0.94 Å on all amino acid residues. Gur-1 and Gur-2 show some differences in binding affinities for T1R2 and T1R3 monomers and the T1R2/T1R3 heterodimer, in addition that only Gur-2 interacts with hT1R2/T1R3 (Table 1). In the aim to explain these differences, molecular dynamics (MD) simulations with Gur-1 and Gur-2 for 100 ns were conducted in GROMACS to compare their stability and conformational changes. As shown in Fig. 6B, Gur-2 has less conformational flexibility than Gur-1. The most flexible part of Gur-1 and Gur-2 is the N-terminal extremity. Interestingly, the two other more flexible regions are those containing Tyr13-Tyr14 and Trp28-Trp29 amino acids of Gur-1. This result is consistent with that of Sigoillot and coworkers who demonstrated the flexibility of the Tyr13-Asp16 amino acid cluster and the Trp28-Trp29 residues. Tyr13-Tyr14 and Trp28-Trp29 are important for the suppressive effect of Gur-1 (Sigoillot et al., 2018), and in this study they are found the principally amino acids interacting with T1R2 and T1R3. Of note, the difference in the flexibility between Gur-1 and Gur-2 is more pronounced in the region containing Trp28-Trp29. To characterize the dynamic changes of the whole peptide, the time evolution of the radius of gyration (Rg) and solvent-accessible surface area (SASA) of Gur-1 and Gur-2 are displayed (Fig. 6C and D). Rg and SASA do not show a great difference between Gur-1 and Gur-2. As mentioned above there is a difference in hydrophobicity, Gur-2 is more hydrophobic than Gur-1 (Table S1). This difference in hydrophobicity, among others, may explain why Gur-2 interacts with human T1R2/T1R3.

**Fig. 6.**
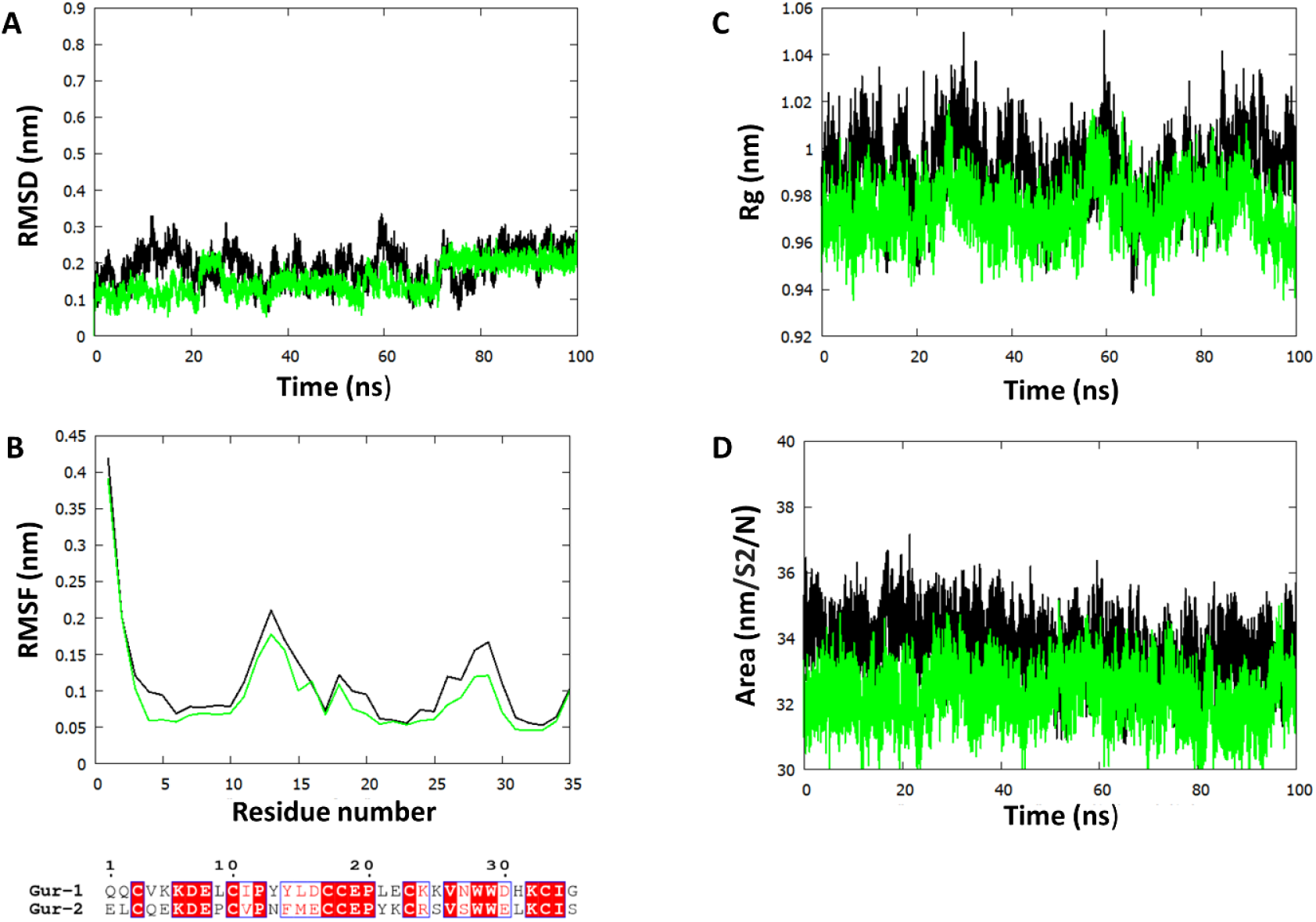
Molecular dynamics simulations (100 ns) of Gur-1 (black curve) and Gur-2 (green curve). **A**, Time evolution of Root mean square deviation (RMSD) with respect to initial structure. **B**, root mean square fluctuations (RMSF) of peptides residues. The position of amino acid residues in alignment is identical to this in RMSF plot. **C** and **D** radius of gyration (Rg) and solvent-accessible surface area (SASA), respectively.

### Interaction of T1R2/T1R3 and Gα-gustducin

On the tongue, the T1R2/T1R3 receptors are coupled with Gα-gustducin for the transmission of the sweet taste signal from taste cell surface to intracellular signaling pathways (Lindemann, 2001; Yasumatsu et al., 2009). Gα-gustducin has been shown to be a key molecule involved in the transduction pathway for Gur-1 sensitive sweet responses (Shigemura et al., 2008; Ohkuri et al., 2009; Yasumatsu et al., 2009).

AF-M predicted that mouse Gα-gustducin (mGα-gustducin) interacts only with the ICD of the mT1R2 subunit in the mT1R2/T1R3 complex (Fig. 7). This result is in agreement with the results of Sainz and coworkers who suggested that the T1R2 subunit in sweet taste receptor is responsible for the activation of the coupled G protein (Sainz et al., 2007), and of Liu and collaborators who showed that heterodimeric metabotropic glutamate receptors (mGlus) transduce signals with one specific subunit that activates G protein (Liu et al., 2017).

**Fig. 7.**
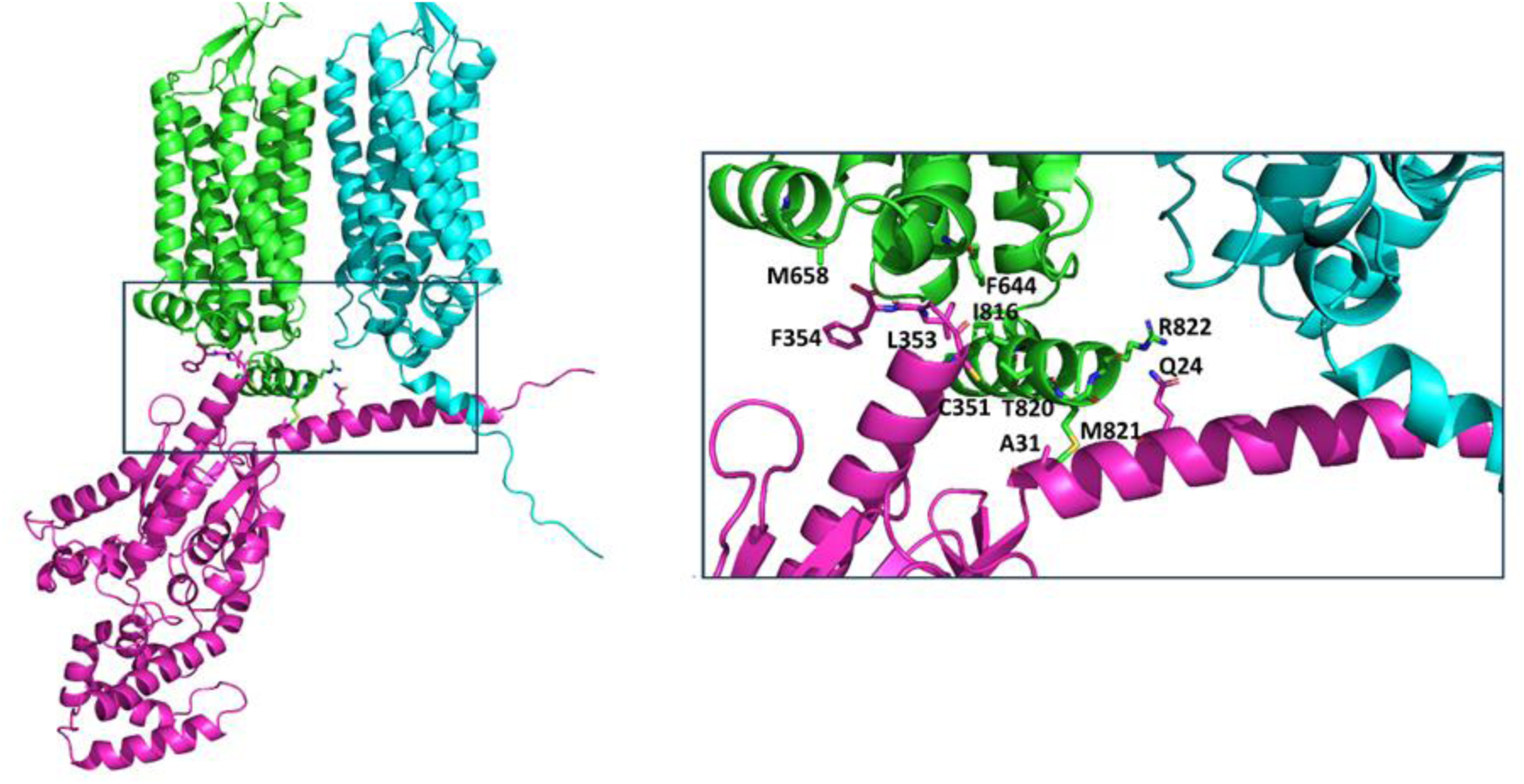
Interaction of Gα-gustducin with the mouse T1R2/T1R3. Gα-gustducin (pink) binds to the intracellular domain (ICD) of T1R2 (green) subunit of T1R2/T1R3 heterodimer (green/cyan). Amino acid residues of T1R2 and T1R3 are numbered without the signal peptide.

In the binding interface, the C-terminus Cys351, Leu353 and Phe354 residues of mGα-gustducin interact with Thr820, Ile816 and Met658 residues of the mT1R2 subunit of the mT1R2/T1R3 complex, respectively. This result is consistent with the work of Hara and collaborators who showed that the last C-terminus 37 amino acid residues of Gα-gustducin are important for its activity in yeast (Hara et al., 2012). Additionally, a point mutation Gly352Pro of Gα-gustducin impaired its ability to activate taste GPCRs (Ruiz-Avila et al., 2001). It is probable that the Gly352Pro mutation destabilizes the region containing residues Cys351, Leu353 and Phe354, preventing Gα-gustducin to interact with the receptor.

## Conclusion

This study sheds light on the discovery of novel gurmarin-like peptides in *G. sylvestre*, in addition to the prediction and the validation of the complete architecture of sweet taste receptor (STR) and the putative binding sites of Gur-1, Gur-2 and Gα-gustducin. It also validates previous hypotheses that suggested that hydrophobic residues Tyr13-Tyr14 and Trp28-Trp29 are involved in the STR-Gur-1 interaction. The putative binding sites will allow to perform the site-directed mutagenesis of specific amino acids in T1R2 and T1R3 to confirm the validity of AF-M predictions. Interestingly, the putative binding site of Gur-2 in the human STR pave the way for improving the binding affinity of Gur-2 by site-directed mutagenesis, and therefore generating peptides with a potent suppressive sweet taste effect in humans. Moreover, the comparison of the binding of Gur-1, Gur-2 and Brz to mT1R2/T1R3 and hT1R2/T1R3 revealed that in Gur-1 and Gur-2 as in Brz, two adjacent aromatic amino acids are important for the interactions with the CRD of T1R2 and T1R3 subunits. It is also interesting to use Gur-1 and Gur-2 (and their isoforms) together in the future experiments to know if they may have a synergistic effect on the suppression of the sweet taste in humans. Finally, gurmarin-like peptides could be interesting drug scaffolds which may help to reduce the craving for sugar and thus help to regulate short-term blood glucose levels.

## Acknowledgments

I thank members of the Centre of Bioinformatics at the Institut de Biologie Intégrative et des Systèmes (IBIS), Université Laval for their assistance.

## Supplementary Material

**Table S1.**
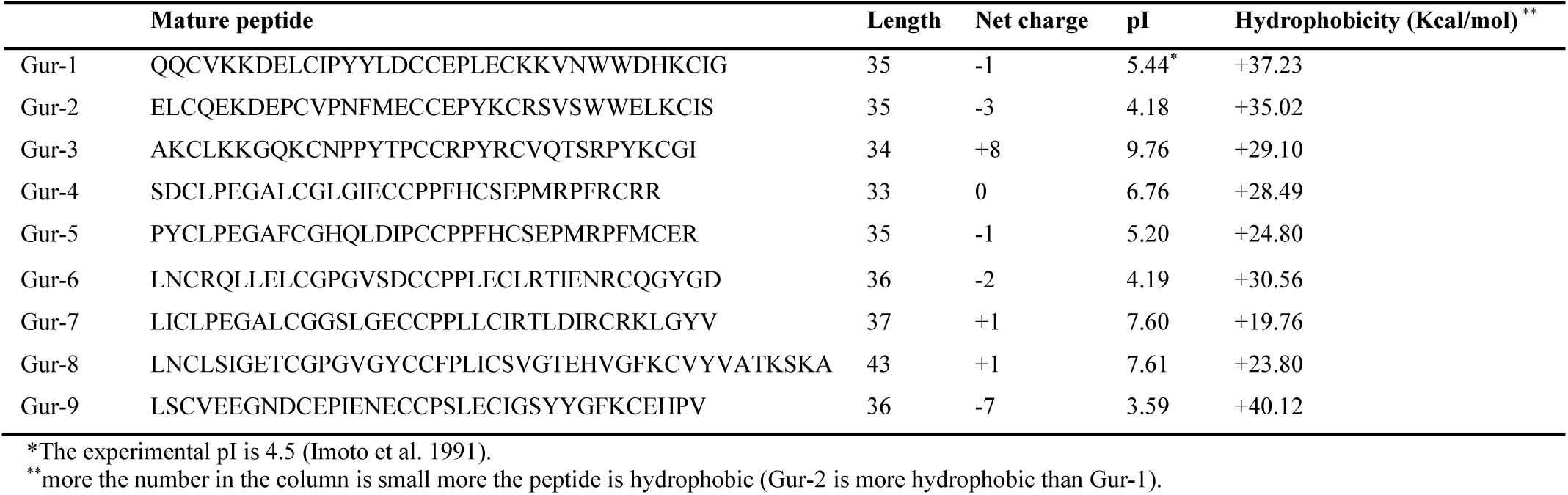
Physico-chemical characteristics of Gur-1 and gurmarin-like peptides.

**Table S2.**
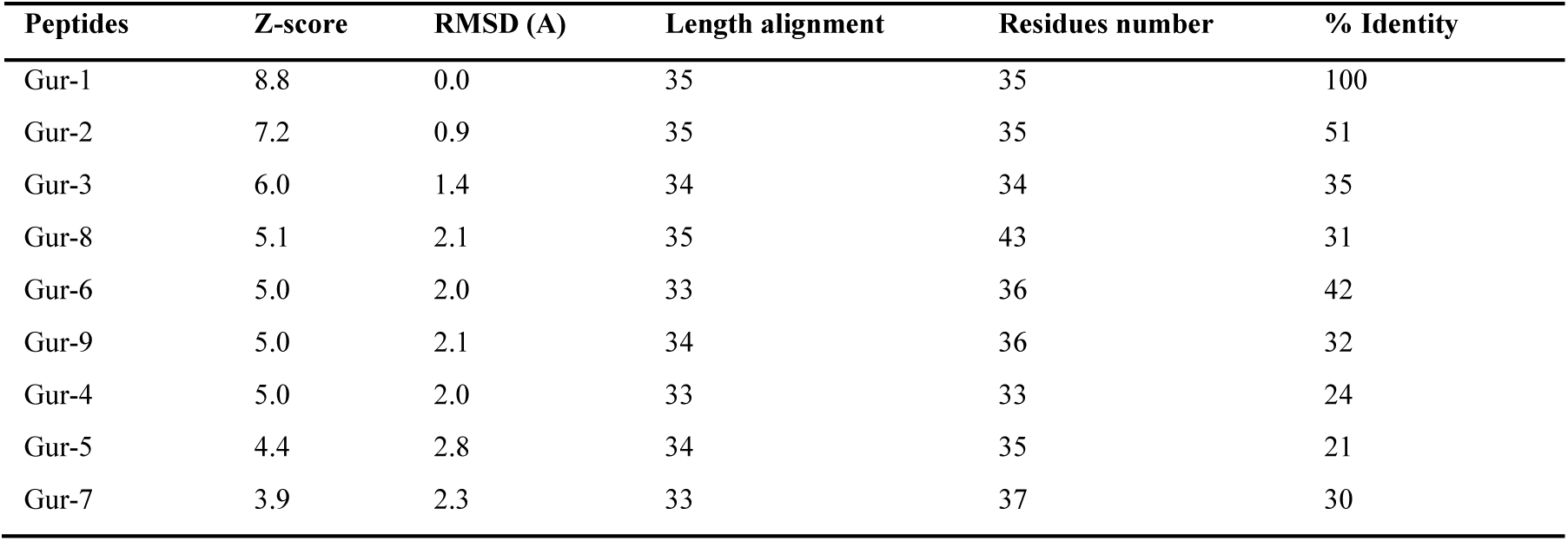
Pairwise structure alignment of Gur-2 to Gur-9 with Gur-1.

**Table S3.**
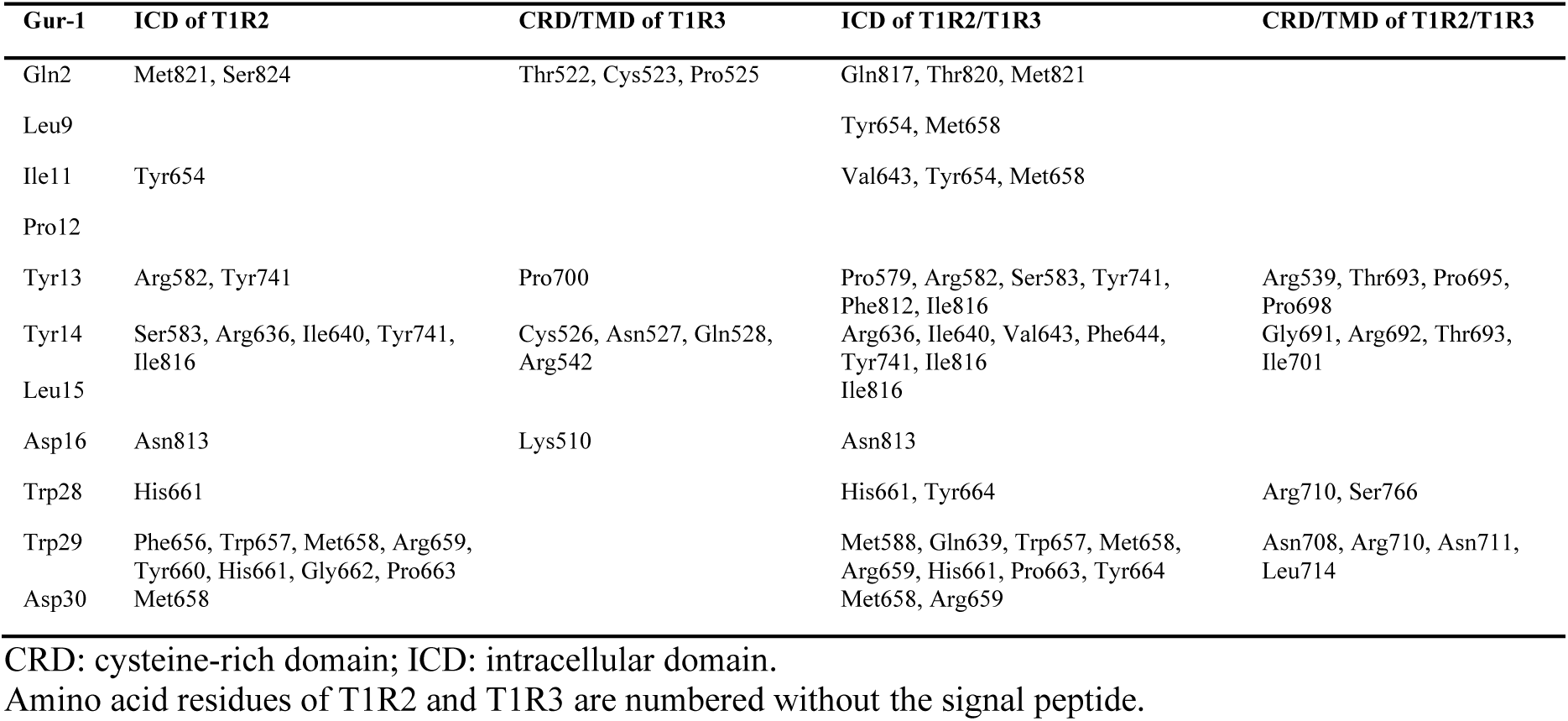
Interactions (≤ 3.5 Å) of Gur-1 amino acid residues with mouse T1R2, T1R3 monomers and T1R2/T1R3 heterodimer.

**Table S4.**
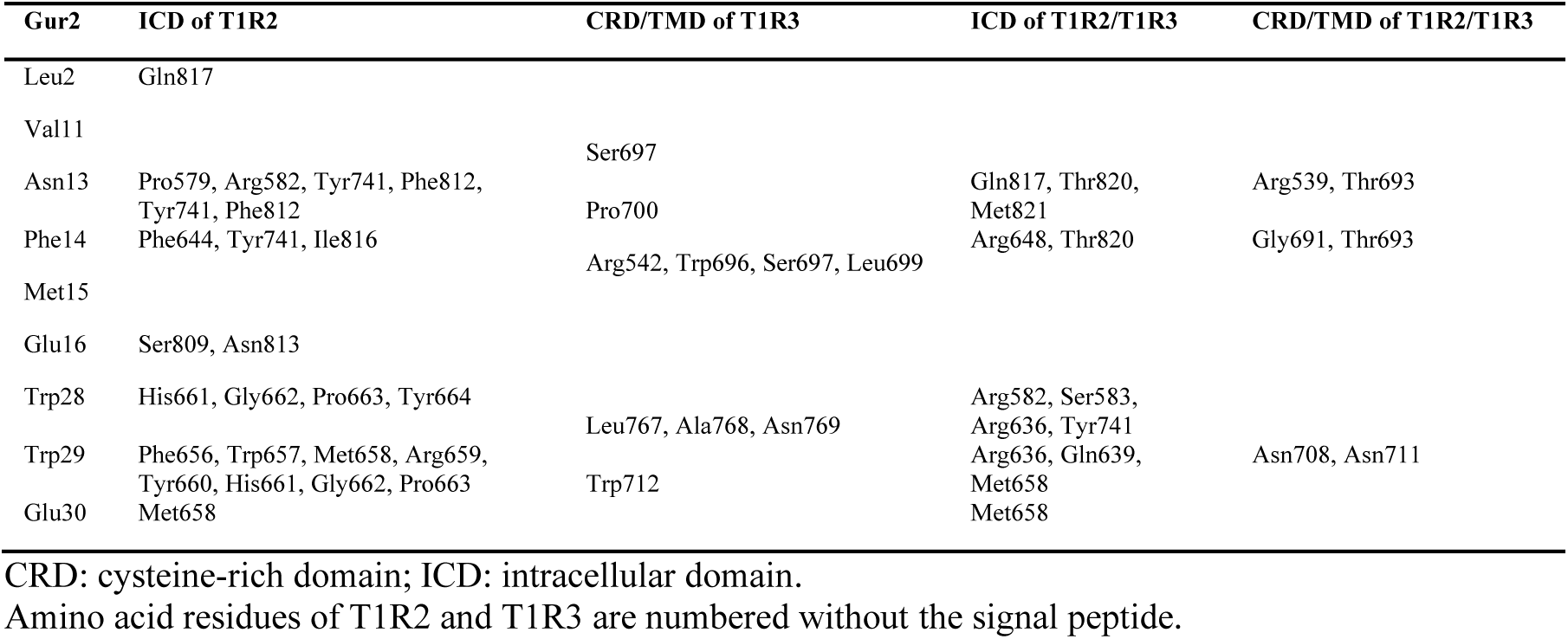
Interactions (≤ 3.5 Å) of Gur-2 amino acid residues with mouse T1R2, T1R3 monomers and T1R2/T1R3 heterodimer.

**Table S5.**
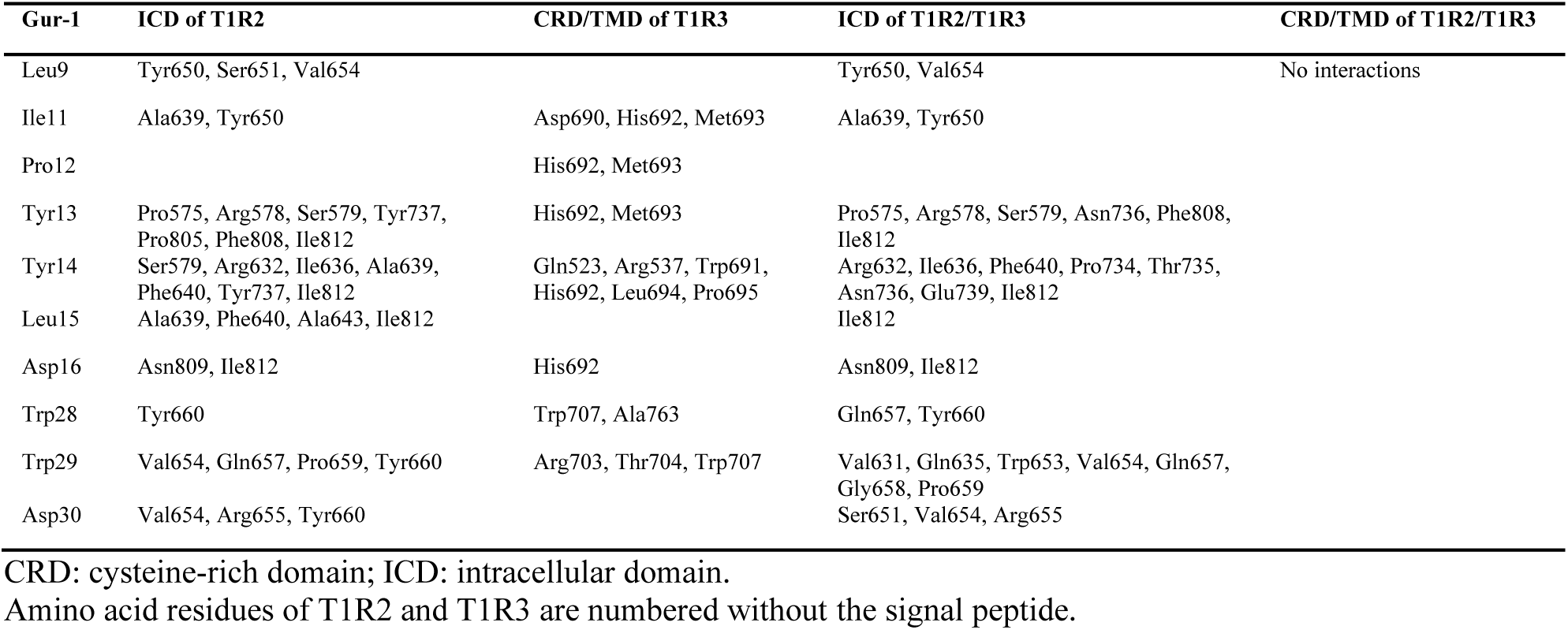
Interactions (≤ 3.5 Å) of Gur-1 amino acid residues with human T1R2, T1R3 monomers and T1R2/T1R3 heterodimer.

**Table S6.**
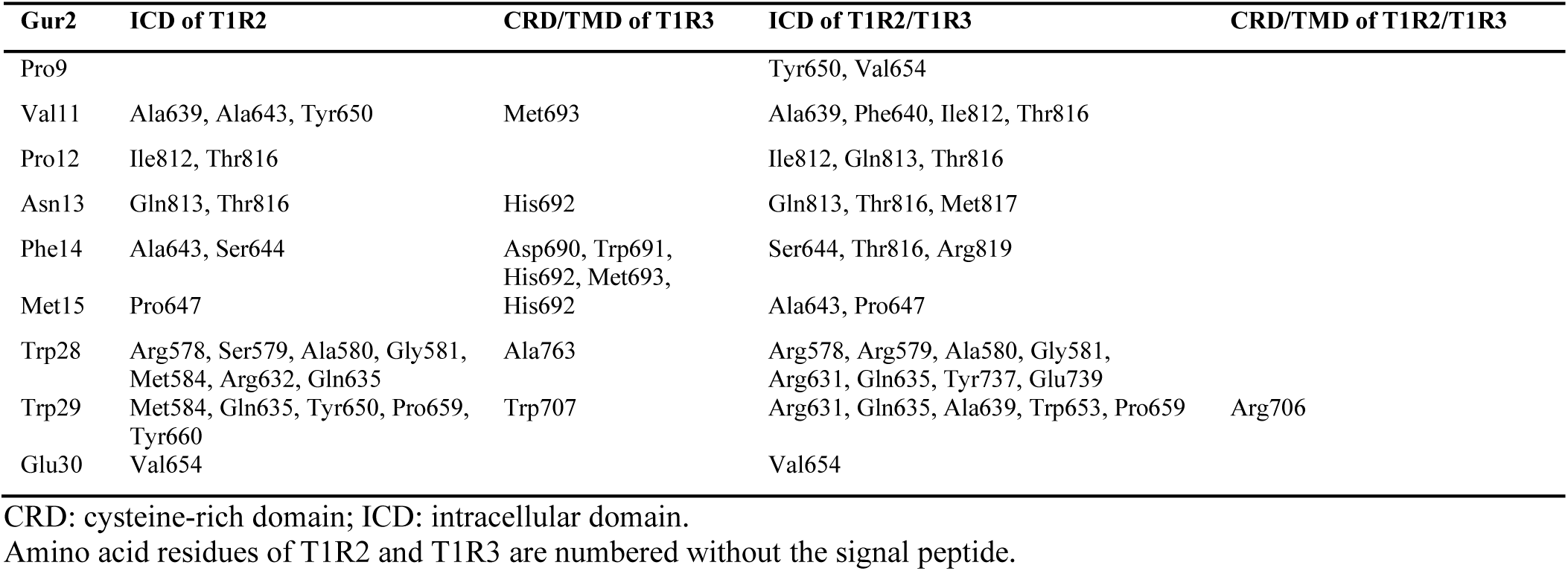
Interactions (≤ 3.5 Å) of Gur-2 amino acid residues with human T1R2, T1R3 monomers and T1R2/T1R3 heterodimer.

**Table S7.**
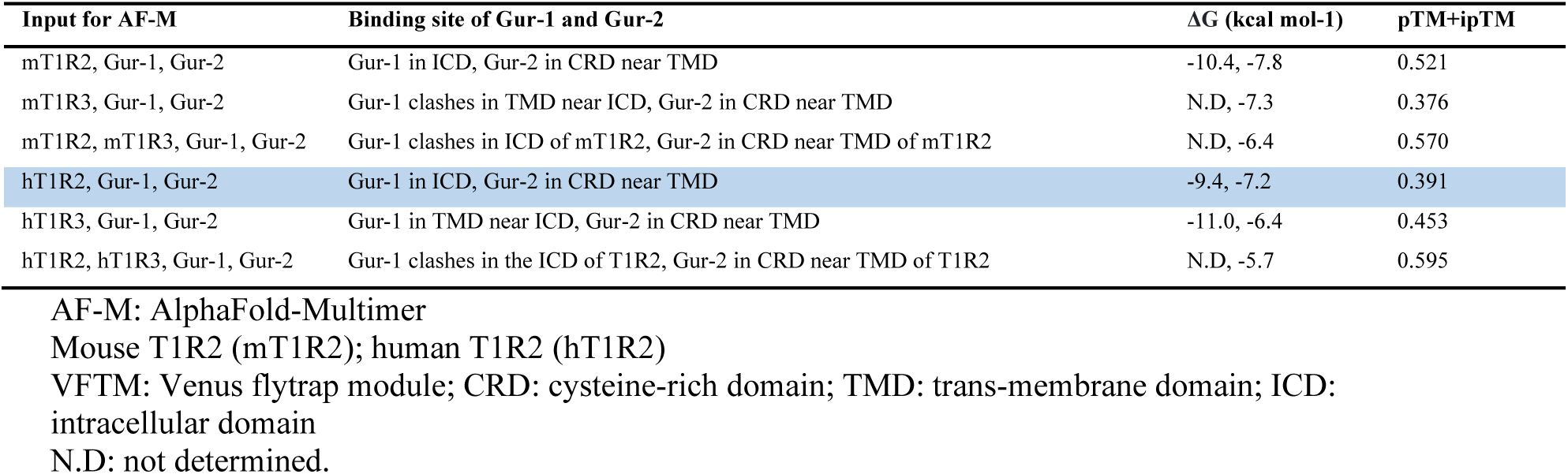
*In silico* assay competitions between Gur-1, Gur-2, T1R2, T1R3 monomers and T1R2/T1R3 heterodimer.

**Fig. S1.**
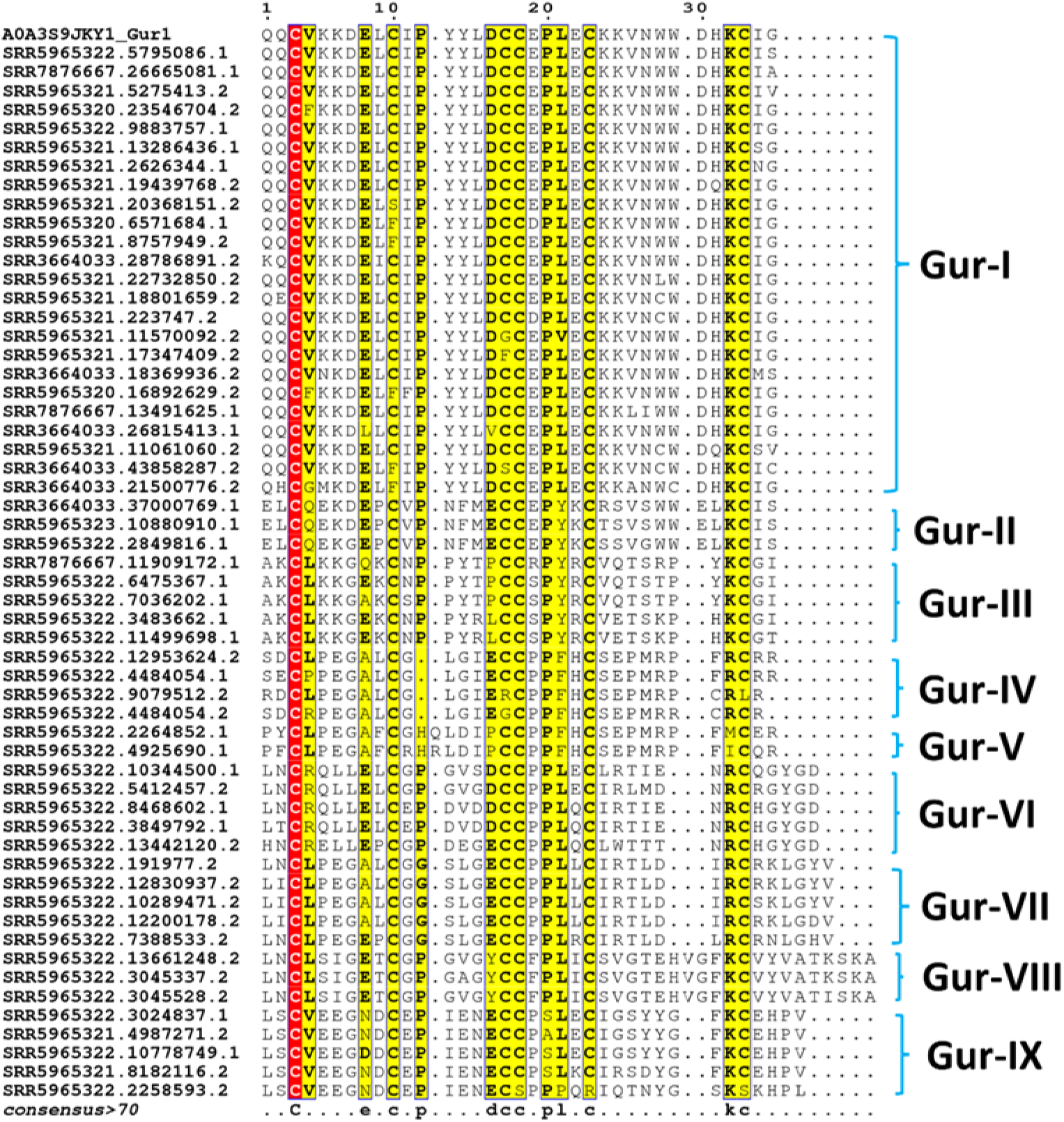
Multiple sequence alignment of *G. sylvestre* gurmarin-like peptides. The aligned sequences are separated in nine groups, named Gur-I to Gur-IX. Gur-I group that contains Gur-1 (UniProt ID: A0A3S9JKY1) and its isoforms is the large group. Trp28 and Trp29 that are important for the suppressive sweet taste effect is conserved in Gur-I and Gur-II groups. SRA accession ID is indicated with each sequence.

**Fig. S2.**
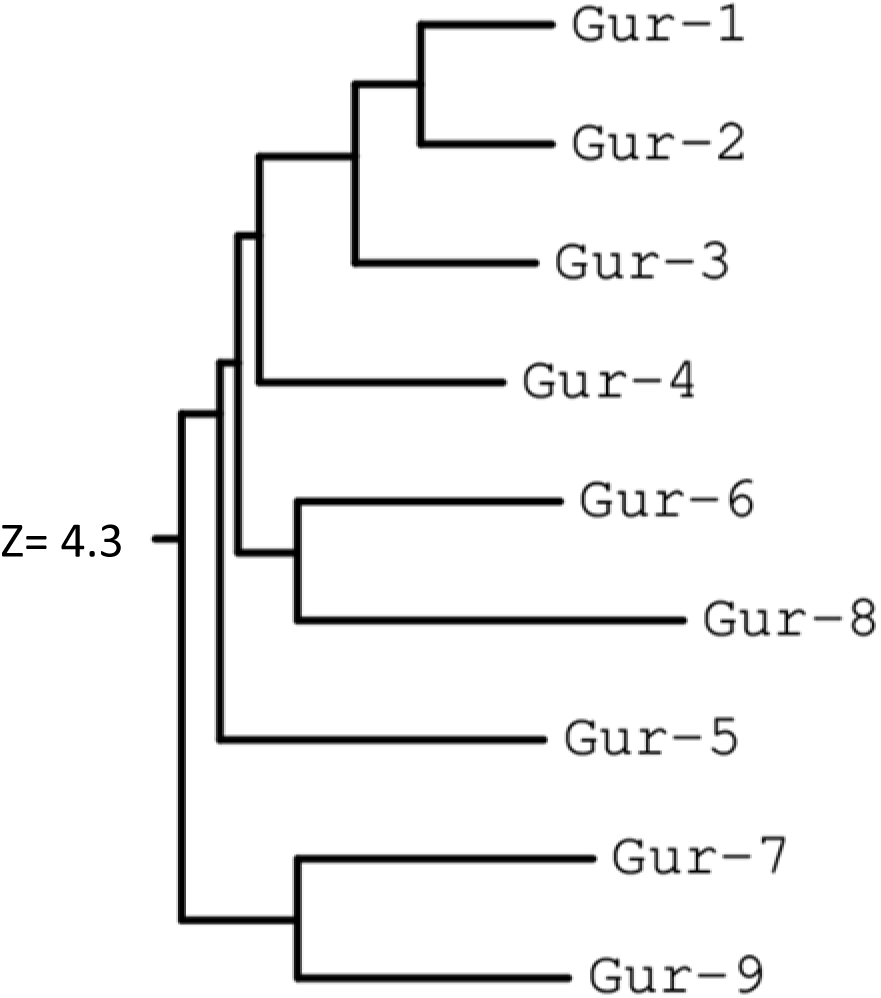
Structural similarity dendrogram of Gur-1 to Gur-9. Labels are linked to structural summaries. The dendrogram is derived by average linkage clustering of the structural similarity matrix (Dali Z-scores, see table S2).

**Fig. S3.**
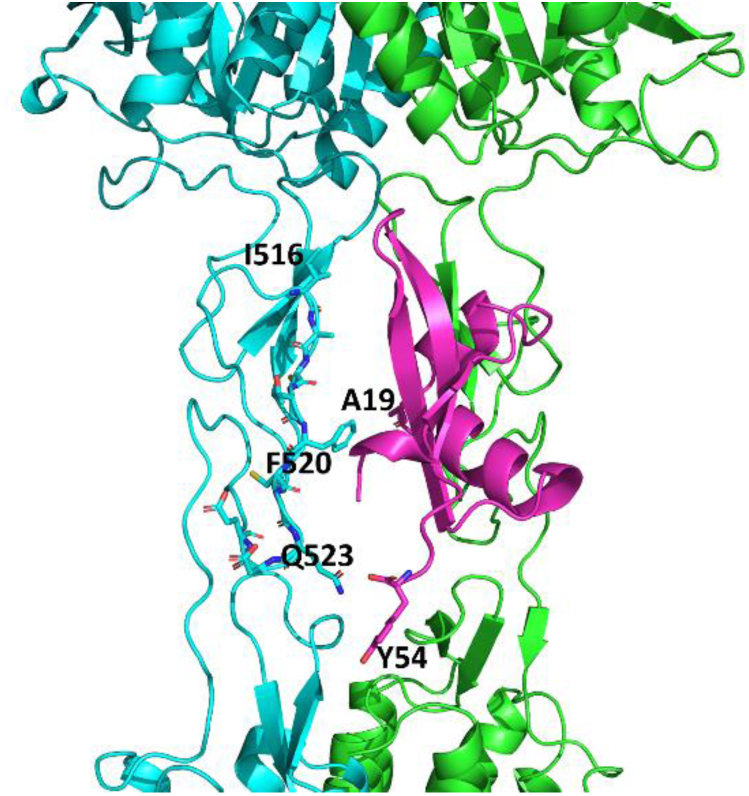
Interactions of Brazzein (Brz) with human T1R2/T1R3 heterodimer. Brz (pink) binds in cysteine-rich domain (CRD, region I516-E525 in sticks) of hT1R3 (cyan) with A19 and Y54 amino acid residues that are important for sweet taste effect of Brz. Amino acid residues of T1R3 are numbered without the signal peptide.

**Fig. S4.**
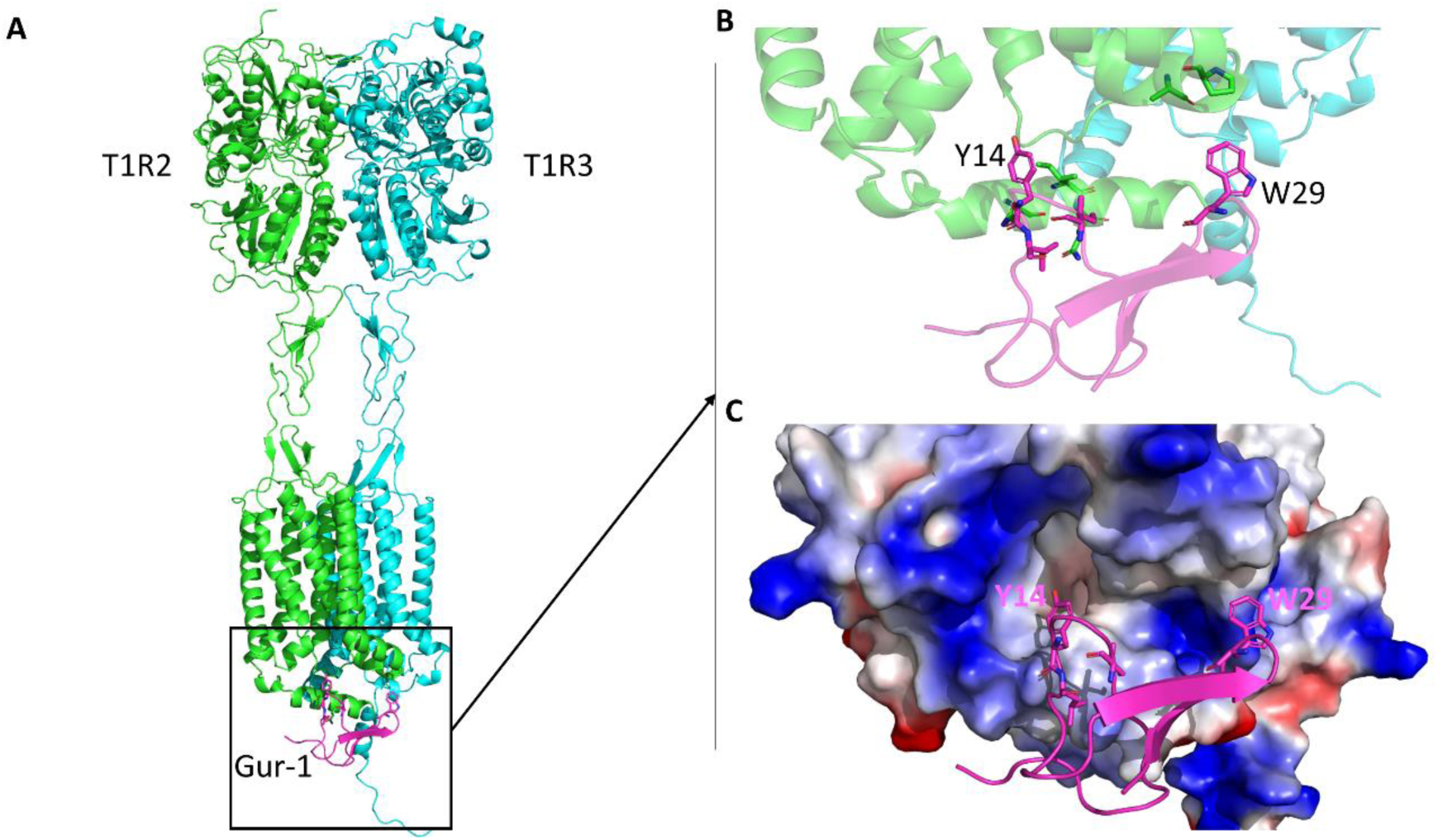
The interaction of Gur-1 with the mouse sweet taste receptor T1R2/T1R3 (m T1R2/T1R3). (**A**) the interaction of Gur-1 (pink) with the intracellular domain (ICD) of mT1R2. (**B**) zoom in the region of the interaction of Gur-1 with the ICD of mT1R2. (**C**) electrostatic potential of the ICD of mT1R2. The negatively, positively and hydrophobic amino acid residues are in red, blue and white, respectively.

**Fig. S5.**
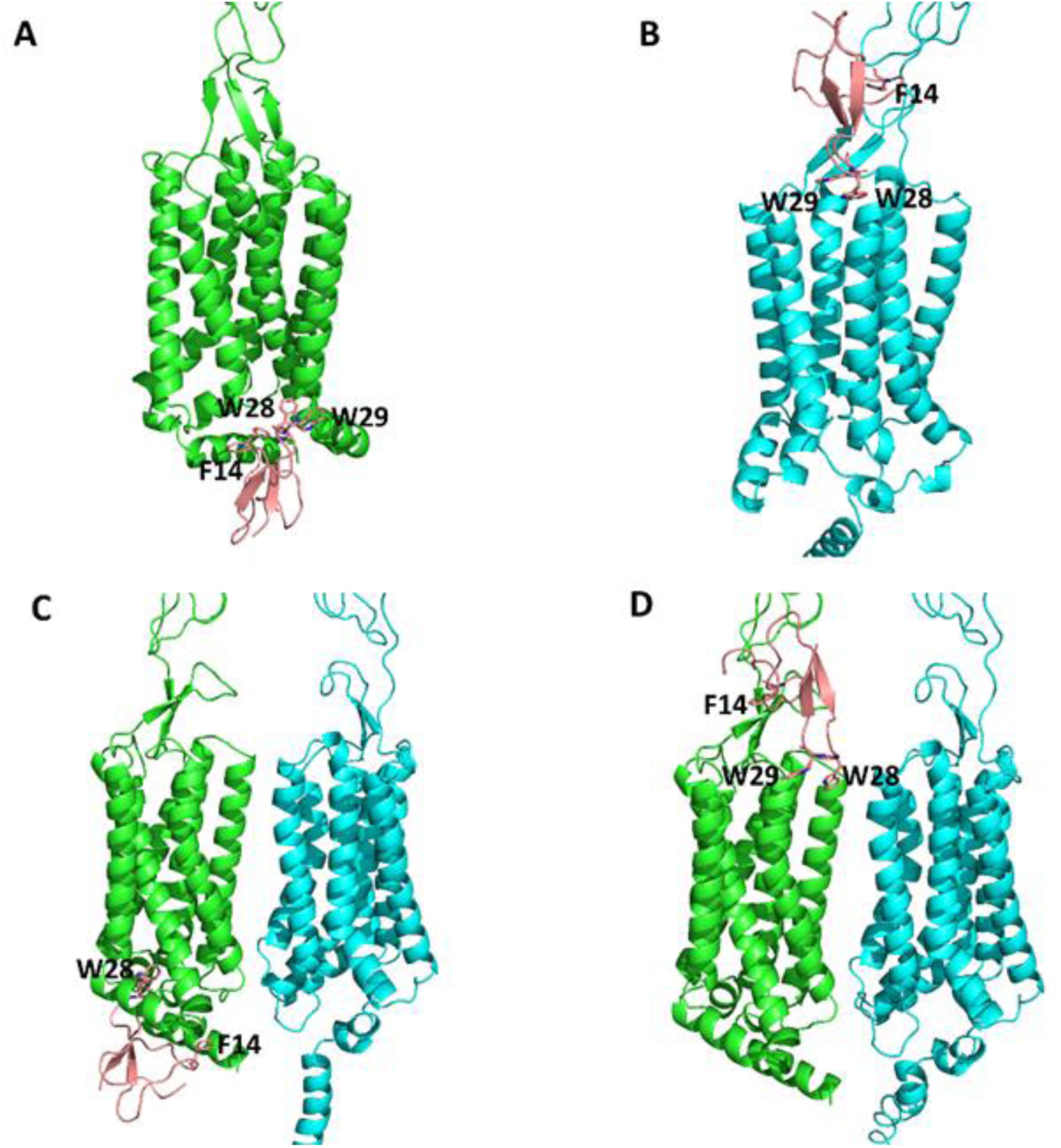
Interactions of Gur-2 with the mouse T1R2, T1R3 monomers and T1R2/T1R3 heterodimer. (**A**) and (**B**) Gur-2 (salmon) binds in the ICD of the T1R2 monomer (green) and in the CRD near TMD of the T1R3 monomer (cyan) with F14, W28 and W29 amino acid residues, respectively. (**C**) and (**D**) Gur-2 binds in the ICD and the CRD near TMD of T1R2 subunit of T1R2/T1R3 heterodimer, respectively. For details of bonds between amino acid residues, see Table S4.

**Fig. S6.**
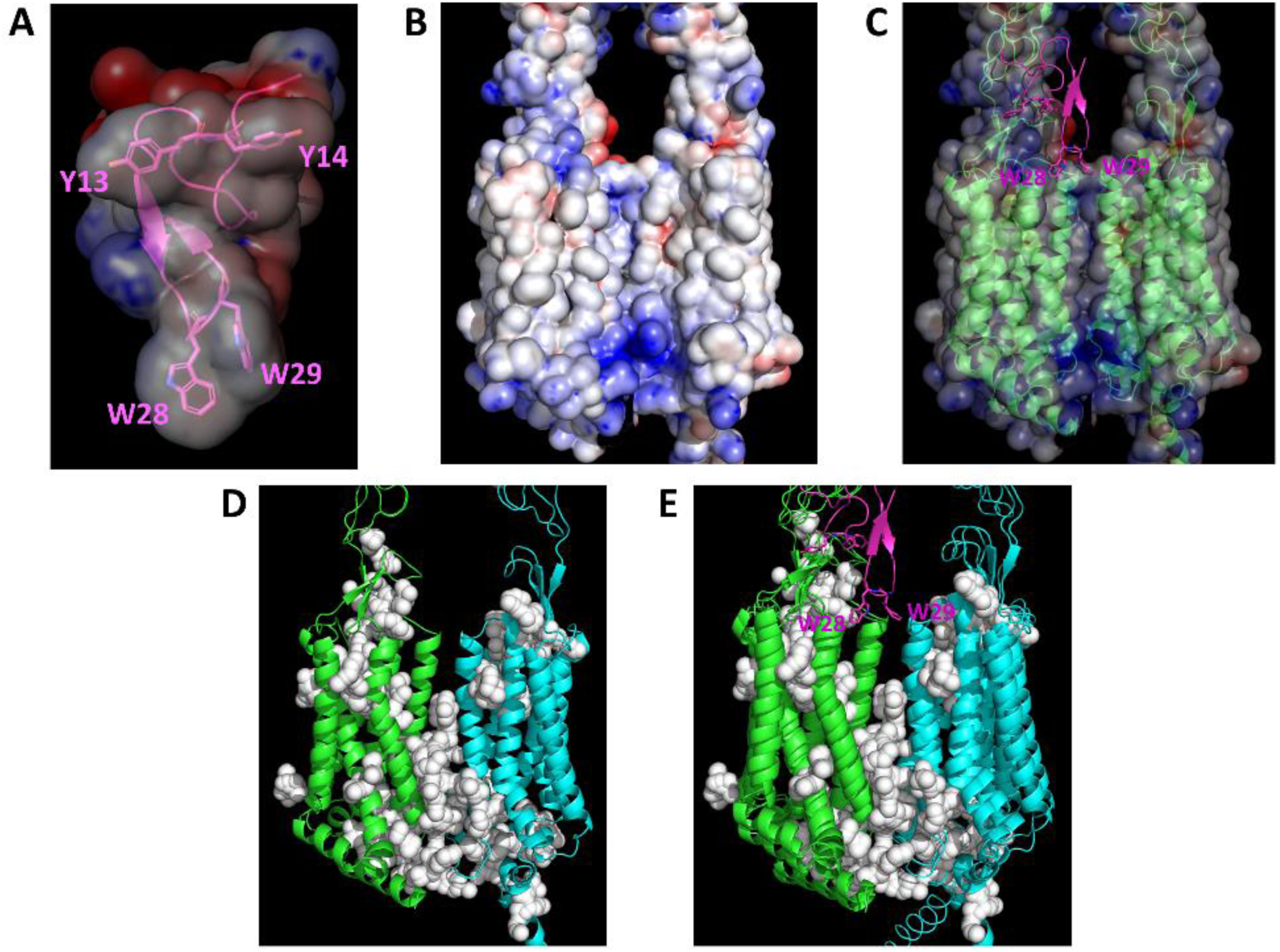
The prediction of tunnels and channels in mouse T1R2/T1R3 structures. (A) electrostatic potential of Gur-1. Important amino acids Y13-14 and W28-29 for sweet suppressive effect are present in hydrophobic region. (B) electrostatic potential of mT1R2/T1R3. The negatively, positively and hydrophobic amino acid residues are in red, blue and white, respectively. (C) superposition of mT1R2/T1R3-Gur-1 complex and electrostatic potential of mT1R2/T1R3. Gur-1 binds in the channel entrance. (D) predicted tunnels and channels in mT1R2-R3 (white spheres). (E) superposition of mT1R2/T1R3-Gur-1 complex and representation of tunnels and channels of mT1R2/T1R3. Gur-1 binds in the channel entrance. T1R2, T1R3 and Gur-1 are in green, cyan and pink, respectively.

## Notes

### Competing Interest Statement

The authors have declared no competing interest.

### Summary of Updates

-All predictions with AF2 and AF-M are performed in triplicate. -Other results are added. -Improved the text.

