## Supplementary Material for "Novel Gurmarin-like Peptides from *Gymnema sylvestre* and their Interactions with the Sweet Taste Receptor T1R2/T1R3"

**Table S1.** Physico-chemical characteristics of Gur-1 and gurmarin-like peptides

| | Mature peptide | Length | Net charge | pI | Hydrophobicity (Kcal/mol) ** |
|---|---|---|---|---|---|
| Gur-1 | QQCVKKDELCIPYYLDCCEPLECKKVNWWDHKCIG | 35 | -1 | 5.44* | +37.23 |
| Gur-2 | ELCQEKDEPCVPNFMECCEPYKCRSVSWWELKCIS | 35 | -3 | 4.18 | +35.02 |
| Gur-3 | AKCLKKGQKCNPPYTPCCRPYRCVQTSRPYKCGI | 34 | +8 | 9.76 | +29.10 |
| Gur-4 | SDCLPEGALCGLGIECCPPFHCSEPMRPFRCRR | 33 | 0 | 6.76 | +28.49 |
| Gur-5 | PYCLPEGAFCGHQLDIPCCPPFHCSEPMRPFMCER | 35 | -1 | 5.20 | +24.80 |
| Gur-6 | LNCRQLLELCGPGVSDCCPPLECLRTIENRCQGYGD | 36 | -2 | 4.19 | +30.56 |
| Gur-7 | LICLPEGALCGGSLGECCPPLLCIRTLDIRCRKLGYV | 37 | +1 | 7.60 | +19.76 |
| Gur-8 | LNCLSIGETCGPGVGYCCFPLICSVGTEHVGFKCVYVATKSKA | 43 | +1 | 7.61 | +23.80 |
| Gur-9 | LSCVEEGNDCEPIENECCPSLECIGSYYGFKCEHPV | 36 | -7 | 3.59 | +40.12 |

*The experimental pI is 4.5 (Imoto et al. 1991).
**more the number in the column is small more the peptide is hydrophobic (Gur-2 is more hydrophobic than Gur-1).

**Table S2**. Pairwise structure alignment of Gur-2 to Gur-9 with Gur-1

| Peptides | Z-score | RMSD (A) | Length alignment | Residues number | % Identity |
|---|---|---|---|---|---|
| Gur-1 | 8.8 | 0.0 | 35 | 35 | 100 |
| Gur-2 | 7.2 | 0.9 | 35 | 35 | 51 |
| Gur-3 | 6.0 | 1.4 | 34 | 34 | 35 |
| Gur-8 | 5.1 | 2.1 | 35 | 43 | 31 |
| Gur-6 | 5.0 | 2.0 | 33 | 36 | 42 |
| Gur-9 | 5.0 | 2.1 | 34 | 36 | 32 |
| Gur-4 | 5.0 | 2.0 | 33 | 33 | 24 |
| Gur-5 | 4.4 | 2.8 | 34 | 35 | 21 |
| Gur-7 | 3.9 | 2.3 | 33 | 37 | 30 |

**Table S3**. Interactions (≤ 3.5 Å) of Gur-1 amino acid residues with mouse T1R2, T1R3 monomers and T1R2/T1R3 heterodimer.

| Gur-1 | ICD of T1R2 | CRD/TMD of T1R3 | ICD of T1R2/T1R3 | CRD/TMD of T1R2/T1R3 |
|---|---|---|---|---|
| Gln2 | Met821, Ser824 | Thr522, Cys523, Pro525 | Gln817, Thr820, Met821 | |
| Leu9 | | | Tyr654, Met658 | |
| Ile11 | Tyr654 | | Val643, Tyr654, Met658 | |
| Pro12 | | | | |
| Tyr13 | Arg582, Tyr741 | Pro700 | Pro579, Arg582, Ser583, Tyr741, Phe812, Ile816 | Arg539, Thr693, Pro695, Pro698 |
| Tyr14 | Ser583, Arg636, Ile640, Tyr741, Ile816 | Cys526, Asn527, Gln528, Arg542 | Arg636, Ile640, Val643, Phe644, Tyr741, Ile816 | Gly691, Arg692, Thr693, Ile701 |
| Leu15 | | | Ile816 | |
| Asp16 | Asn813 | Lys510 | Asn813 | |
| Trp28 | His661 | | His661, Tyr664 | Arg710, Ser766 |
| Trp29 | Phe656, Trp657, Met658, Arg659, Tyr660, His661, Gly662, Pro663 | | Met588, Gln639, Trp657, Met658, Arg659, His661, Pro663, Tyr664 | Asn708, Arg710, Asn711, Leu714 |
| Asp30 | Met658 | | Met658, Arg659 | |

CRD: cysteine-rich domain; ICD: intracellular domain.
Amino acid residues of T1R2 and T1R3 are numbered without the signal peptide.

**Table S4**. Interactions (≤ 3.5 Å) of Gur-2 amino acid residues with mouse T1R2, T1R3 monomers and T1R2/T1R3 heterodimer.

| Gur2 | ICD of T1R2 | CRD/TMD of T1R3 | ICD of T1R2/T1R3 | CRD/TMD of T1R2/T1R3 |
|---|---|---|---|---|
| Leu2 | Gln817 | | | |
| Val11 | | Ser697 | | |
| Asn13 | Pro579, Arg582, Tyr741, Phe812, Tyr741, Phe812 | Pro700 | Gln817, Thr820, Met821 | Arg539, Thr693 |
| Phe14 | Phe644, Tyr741, Ile816 | Arg542, Trp696, Ser697, Leu699 | Arg648, Thr820 | Gly691, Thr693 |
| Met15 | | | | |
| Glu16 | Ser809, Asn813 | | | |
| Trp28 | His661, Gly662, Pro663, Tyr664 | Leu767, Ala768, Asn769 | Arg582, Ser583, Arg636, Tyr741 | |
| Trp29 | Phe656, Trp657, Met658, Arg659, Tyr660, His661, Gly662, Pro663 | Trp712 | Arg636, Gln639, Met658 | Asn708, Asn711 |
| Glu30 | Met658 | | Met658 | |

CRD: cysteine-rich domain; ICD: intracellular domain.
Amino acid residues of T1R2 and T1R3 are numbered without the signal peptide.

**Table S5**. Interactions (≤ 3.5 Å) of Gur-1 amino acid residues with human T1R2, T1R3 monomers and T1R2/T1R3 heterodimer.

| Gur-1 | ICD of T1R2 | CRD/TMD of T1R3 | ICD of T1R2/T1R3 | CRD/TMD of T1R2/T1R3 |
|---|---|---|---|---|
| Leu9 | Tyr650, Ser651, Val654 | | Tyr650, Val654 | No interactions |
| Ile11 | Ala639, Tyr650 | Asp690, His692, Met693 | Ala639, Tyr650 | |
| Pro12 | | His692, Met693 | | |
| Tyr13 | Pro575, Arg578, Ser579, Tyr737, Pro805, Phe808, Ile812 | His692, Met693 | Pro575, Arg578, Ser579, Asn736, Phe808, Ile812 | |
| Tyr14 | Ser579, Arg632, Ile636, Ala639, Phe640, Tyr737, Ile812 | Gln523, Arg537, Trp691, His692, Leu694, Pro695 | Arg632, Ile636, Phe640, Pro734, Thr735, Asn736, Glu739, Ile812 | |
| Leu15 | Ala639, Phe640, Ala643, Ile812 | | Ile812 | |
| Asp16 | Asn809, Ile812 | His692 | Asn809, Ile812 | |
| Trp28 | Tyr660 | Trp707, Ala763 | Gln657, Tyr660 | |
| Trp29 | Val654, Gln657, Pro659, Tyr660 | Arg703, Thr704, Trp707 | Val631, Gln635, Trp653, Val654, Gln657, Gly658, Pro659 | |
| Asp30 | Val654, Arg655, Tyr660 | | Ser651, Val654, Arg655 | |

CRD: cysteine-rich domain; ICD: intracellular domain.
Amino acid residues of T1R2 and T1R3 are numbered without the signal peptide.

**Table S6**. Interactions (≤ 3.5 Å) of Gur-2 amino acid residues with human T1R2, T1R3 monomers and T1R2/T1R3 heterodimer.

| Gur2 | ICD of T1R2 | CRD/TMD of T1R3 | ICD of T1R2/T1R3 | CRD/TMD of T1R2/T1R3 |
|---|---|---|---|---|
| Pro9 | | | Tyr650, Val654 | |
| Val11 | Ala639, Ala643, Tyr650 | Met693 | Ala639, Phe640, Ile812, Thr816 | |
| Pro12 | Ile812, Thr816 | | Ile812, Gln813, Thr816 | |
| Asn13 | Gln813, Thr816 | His692 | Gln813, Thr816, Met817 | |
| Phe14 | Ala643, Ser644 | Asp690, Trp691, His692, Met693, | Ser644, Thr816, Arg819 | |
| Met15 | Pro647 | His692 | Ala643, Pro647 | |
| Trp28 | Arg578, Ser579, Ala580, Gly581, Met584, Arg632, Gln635 | Ala763 | Arg578, Arg579, Ala580, Gly581, Arg631, Gln635, Tyr737, Glu739 | |
| Trp29 | Met584, Gln635, Tyr650, Pro659, Tyr660 | Trp707 | Arg631, Gln635, Ala639, Trp653, Pro659 | Arg706 |
| Glu30 | Val654 | | Val654 | |

CRD: cysteine-rich domain; ICD: intracellular domain.
Amino acid residues of T1R2 and T1R3 are numbered without the signal peptide.

**Table S7**. *In silico* assay competitions between Gur-1, Gur-2, T1R2, T1R3 monomers and T1R2/T1R3 heterodimer.

| Input for AF-M | Binding site of Gur-1 and Gur-2 | ΔG (kcal mol-1) | pTM+ipTM |
|---|---|---|---|
| mT1R2, Gur-1, Gur-2 | Gur-1 in ICD, Gur-2 in CRD near TMD | -10.4, -7.8 | 0.521 |
| mT1R3, Gur-1, Gur-2 | Gur-1 clashes in TMD near ICD, Gur-2 in CRD near TMD | N.D, -7.3 | 0.376 |
| mT1R2, mT1R3, Gur-1, Gur-2 | Gur-1 clashes in ICD of mT1R2, Gur-2 in CRD near TMD of mT1R2 | N.D, -6.4 | 0.570 |
| hT1R2, Gur-1, Gur-2 | Gur-1 in ICD, Gur-2 in CRD near TMD | -9.4, -7.2 | 0.391 |
| hT1R3, Gur-1, Gur-2 | Gur-1 in TMD near ICD, Gur-2 in CRD near TMD | -11.0, -6.4 | 0.453 |
| hT1R2, hT1R3, Gur-1, Gur-2 | Gur-1 clashes in the ICD of T1R2, Gur-2 in CRD near TMD of T1R2 | N.D, -5.7 | 0.595 |

AF-M: AlphaFold-Multimer
Mouse T1R2 (mT1R2); human T1R2 (hT1R2)
VFTM: Venus flytrap module; CRD: cysteine-rich domain; TMD: trans-membrane domain; ICD: intracellular domain
N.D: not determined.

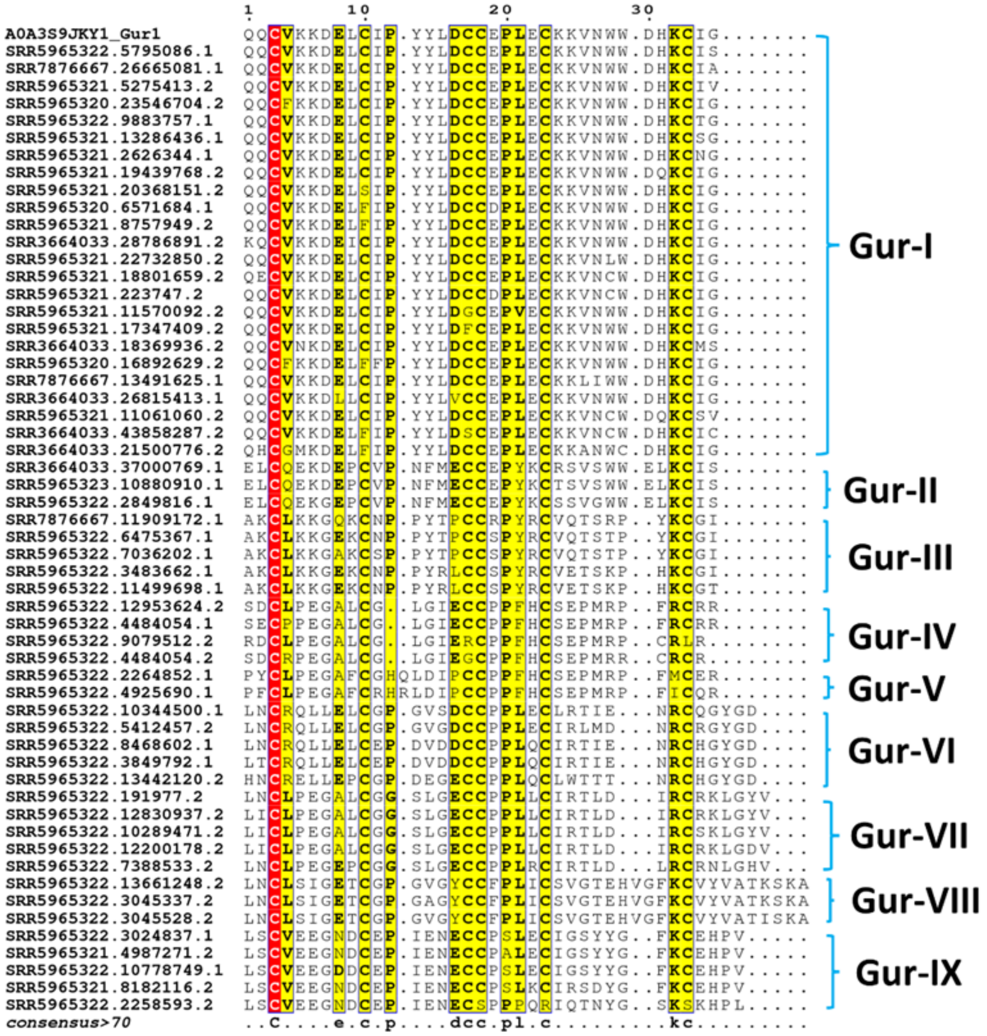


**Fig. S1**. Multiple sequence alignment of *G. sylvestre* gurmarin-like peptides. The aligned sequences are separated in nine groups, named Gur-I to Gur-IX. Gur-I group that contains Gur-1 (UniProt ID: A0A3S9JKY1) and its isoforms is the large group. Trp28 and Trp29 that are important for the suppressive sweet taste effect is conserved in Gur-I and Gur-II groups. SRA accession ID is indicated with each sequence.

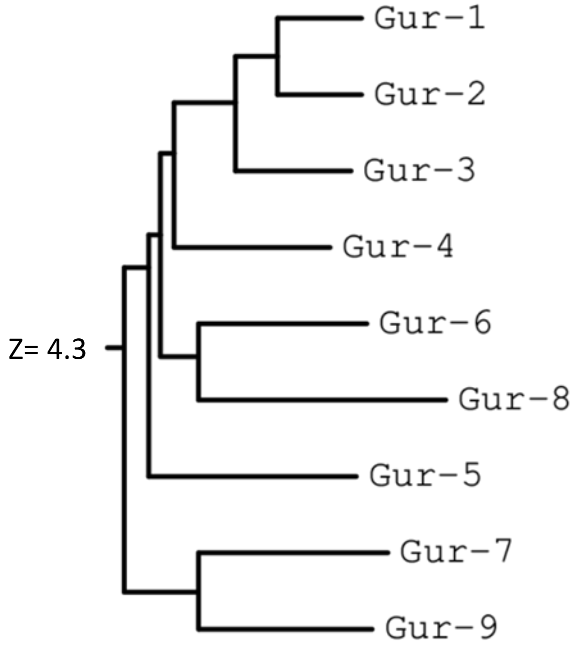


**Fig. S2**. Structural similarity dendrogram of Gur-1 to Gur-9. Labels are linked to structural summaries. The dendrogram is derived by average linkage clustering of the structural similarity matrix (Dali Z-scores, see table S2).

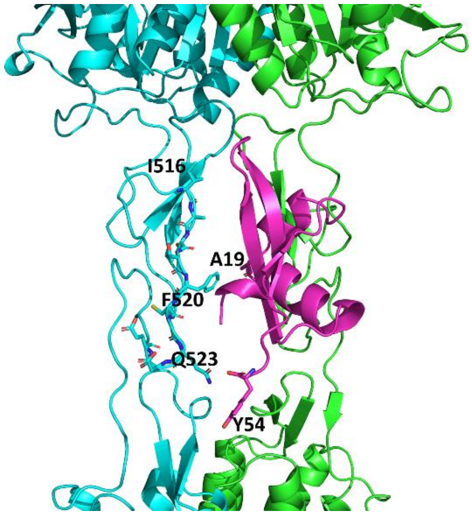


**Fig. S3.** Interactions of Brazzein (Brz) with human T1R2/T1R3 heterodimer. Brz (pink) binds in cysteine-rich domain (CRD, region I516-E525 in sticks) of hT1R3 (cyan) with A19 and Y54 amino acid residues that are important for sweet taste effect of Brz. Amino acid residues of T1R3 are numbered without the signal peptide.

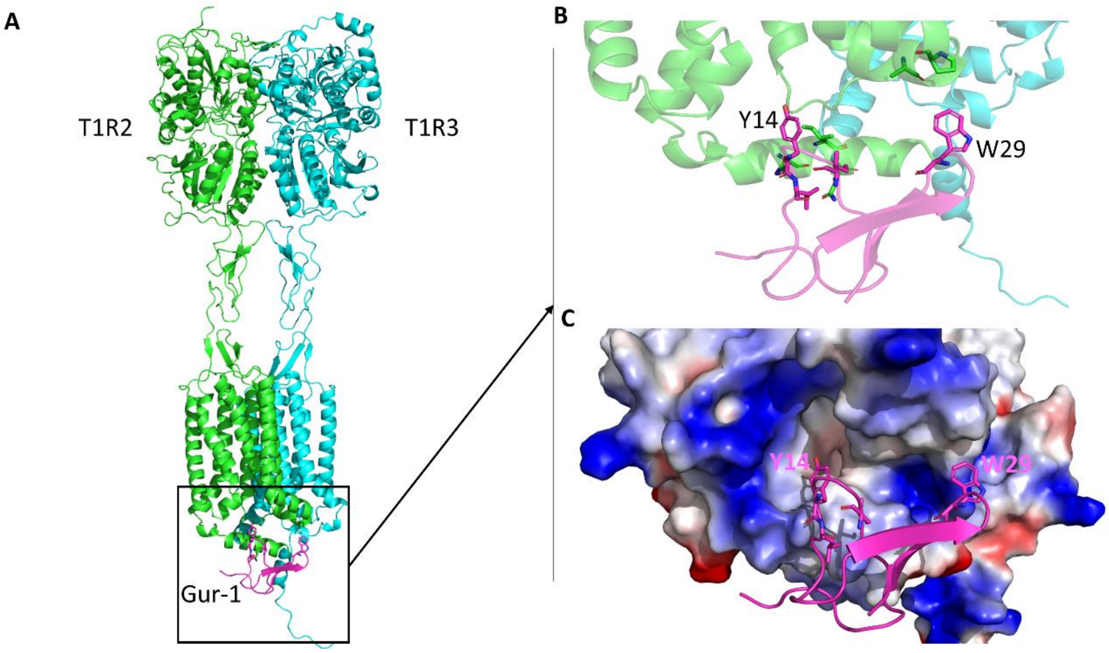


**Fig. S4.** The interaction of Gur-1 with the mouse sweet taste receptor T1R2/T1R3 (m T1R2/T1R3). (**A**) the interaction of Gur-1 (pink) with the intracellular domain (ICD) of mT1R2. (**B**) zoom in the region of the interaction of Gur-1 with the ICD of mT1R2. (**C**) electrostatic potential of the ICD of mT1R2. The negatively, positively and hydrophobic amino acid residues are in red, blue and white, respectively.

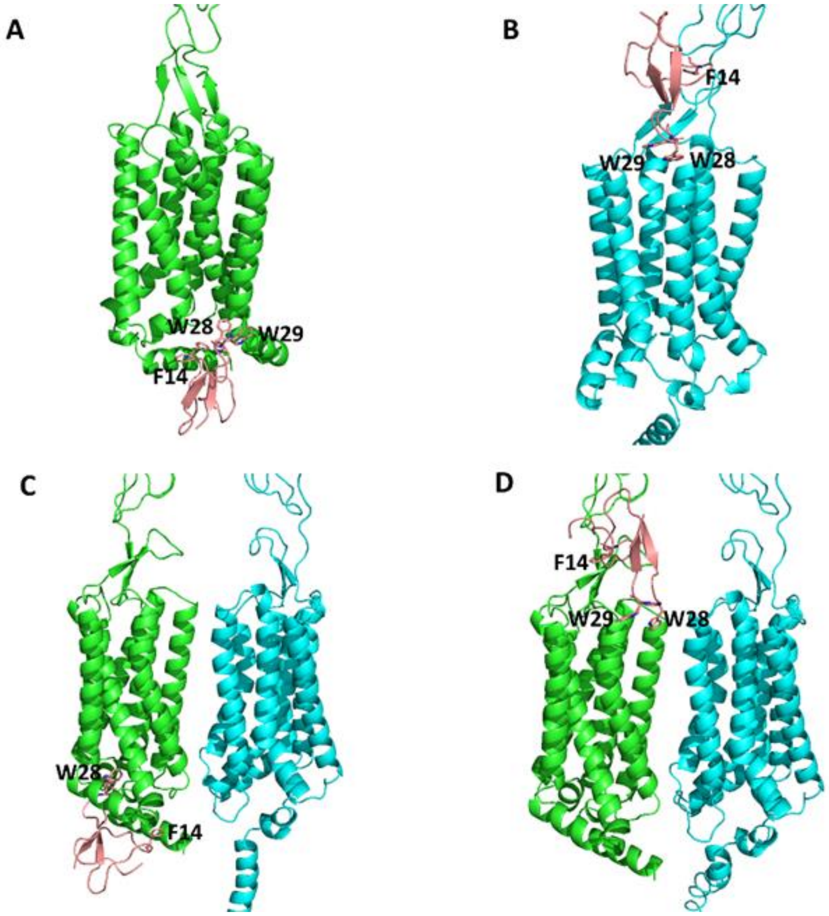


**Fig. S5.** Interactions of Gur-2 with the mouse T1R2, T1R3 monomers and T1R2/T1R3 heterodimer. (**A**) and (**B**) Gur-2 (salmon) binds in the ICD of the T1R2 monomer (green) and in the CRD near TMD of the T1R3 monomer (cyan) with F14, W28 and W29 amino acid residues, respectively. (**C**) and (**D**) Gur-2 binds in the ICD and the CRD near TMD of T1R2 subunit of T1R2/T1R3 heterodimer, respectively. For details of bonds between amino acid residues, see Table S4.

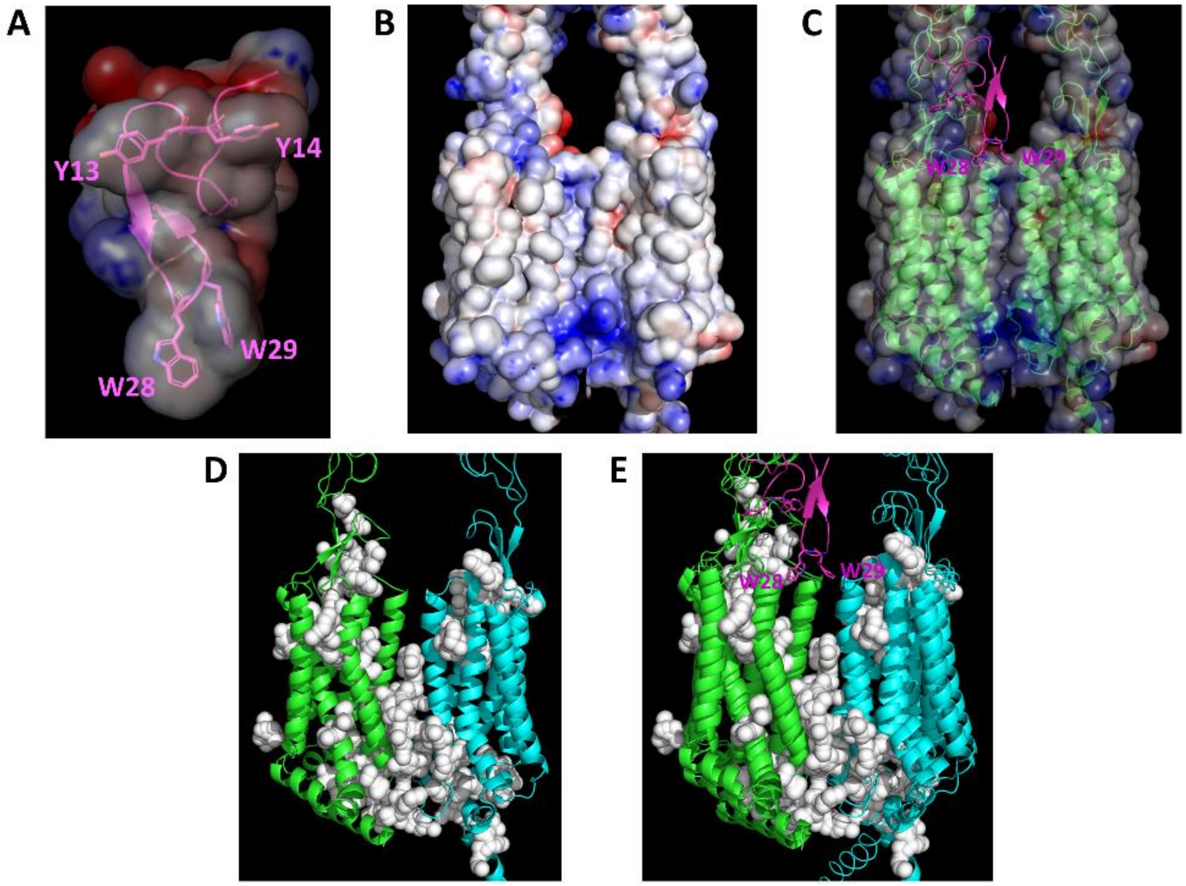


**Fig. S6.** The prediction of tunnels and channels in mouse T1R2/T1R3 structures. (A) electrostatic potential of Gur-1. Important amino acids Y13-14 and W28-29 for sweet suppressive effect are present in hydrophobic region. (B) electrostatic potential of mT1R2/T1R3. The negatively, positively and hydrophobic amino acid residues are in red, blue and white, respectively. (C) superposition of mT1R2/T1R3-Gur-1 complex and electrostatic potential of mT1R2/T1R3. Gur-1 binds in the channel entrance. (D) predicted tunnels and channels in mT1R2-R3 (white spheres). (E) superposition of mT1R2/T1R3-Gur-1 complex and representation of tunnels and channels of mT1R2/T1R3. Gur-1 binds in the channel entrance. T1R2, T1R3 and Gur-1 are in green, cyan and pink, respectively.
